# Convergent suppression of nuclear-encoded mitochondrial fatty acid oxidation genes defines a pan-subtype signature in breast cancer: a multi-cohort transcriptomic study

**DOI:** 10.64898/2026.05.17.725700

**Authors:** Aroob Gomosani, Haneen Marghalani, Layal Al Matar

## Abstract

**Background:** Breast cancer exhibits extensive molecular heterogeneity across intrinsic subtypes, yet convergent metabolic reprogramming may represent an obligate feature of tumour initiation. We hypothesised that suppression of nuclear-encoded mitochondrial fatty acid oxidation (FAO) constitutes such a convergence point, defining a shared metabolic phenotype independent of subtype.

**Methods:** RNA-seq data from 1,106 primary breast tumours and 113 normal-adjacent tissues (TCGA-BRCA) were intersected with 1,079 nuclear-encoded mitochondrial genes from MitoCarta 3.0. Differential expression was assessed using Welch t-test with Benjamini–Hochberg correction at all tumour stages, at Stage I specifically, and stratified across PAM50 subtypes. A 55-gene core FAO signature was derived by three-way intersection. Ten candidate genes were selected by pre-specified objective scoring, locked before any clinical testing. Gene set enrichment analysis (GSEA) was performed using MitoCarta 3.0 pathway annotations. Diagnostic performance, clinical associations, survival, and mutation independence were characterised. External validation used two independent GEO cohorts (GSE42568, n = 121; GSE109169, n = 50); prognostic validation used METABRIC (Molecular Taxonomy of Breast Cancer International Consortium; n = 1,980). DESeq2 was applied as methodological cross-validation.

**Results:** Among 126 differentially expressed mitochondrial genes, fatty acid oxidation was the most significantly depleted pathway (normalised enrichment score −2.130; false discovery rate 0.001). The 55-gene core signature replicated in both external cohorts with 100% directional concordance (hypergeometric p < 10⁻¹⁵). All 10 candidate genes discriminated tumour from normal tissue (area under the curve 0.915–0.979) and demonstrated broad clinical associations. The composite FAO suppression score predicted overall survival in METABRIC (log-rank p = 7.82 × 10⁻⁷) and MAOA achieved independent prognostic significance in multivariable Cox regression (hazard ratio 0.890; adjusted p = 0.009). DESeq2 cross-validation confirmed Spearman ρ = 0.980 concordance.

**Conclusions:** Nuclear-encoded FAO suppression is a robust, pan-subtype feature of breast cancer detectable at Stage I and validated across independent platforms and cohorts. These 10 candidate genes constitute a consistent initiation-phase mitochondrial signature, implicating FAO suppression as a potential convergence point in breast cancer oncogenesis and motivating targeted functional investigation.

## Background

Breast cancer is the most commonly diagnosed malignancy in women worldwide, with an estimated 2.3 million new cases annually [1]. Despite transformative advances in molecular subtyping — most notably the PAM50 classifier defining Luminal A, Luminal B, HER2-enriched, and Basal-like intrinsic subtypes [2, 3] — the relationship between subtype-specific oncogenic programmes and shared metabolic reprogramming remains incompletely characterised. Effective therapeutic targeting requires understanding not only what distinguishes molecular subtypes, but what mechanistically unites them.

Metabolic reprogramming is a recognised hallmark of cancer [4]. The preferential utilisation of aerobic glycolysis — the Warburg effect — has been extensively characterised across tumour types [5, 6]. However, the mitochondrial dimension of this metabolic shift, and in particular the status of fatty acid oxidation (FAO), has received comparatively less systematic attention in breast cancer. FAO, which occurs in both mitochondria and peroxisomes and is encoded entirely by nuclear genes, generates acetyl-CoA, NADH, and FADH₂ to support bioenergetic and biosynthetic demands. Its suppression has been proposed to redirect lipid substrates toward anabolic pathways sustaining rapid proliferation [7, 8]. Although subtype-specific alterations in FAO-related enzymes have been reported [9], whether FAO suppression constitutes a convergent, pan-subtype feature present at the earliest detectable disease stage has not been systematically evaluated at genome-wide scale.

The identification of FAO suppression as a potential convergence point in breast oncogenesis would reframe mitochondrial metabolic reprogramming from a downstream consequence of somatic mutation to a concurrent and possibly enabling feature of malignant transformation. Such a reframing implies that oncogenic permissiveness may require a concurrent mitochondrial context in which suppression of oxidative lipid metabolism creates a metabolic environment conducive to tumour initiation. A pan-subtype shared molecular vulnerability would support development of FAO-targeted interventions agnostic to molecular classification — addressing a fundamental limitation of current subtype-stratified treatment paradigms.

We therefore undertook a systematic transcriptomic analysis of the nuclear-encoded mitochondrial proteome in breast cancer. Using TCGA-BRCA as the primary discovery cohort, MitoCarta 3.0 as the definitive mitochondrial gene reference [10], and two independent external cohorts spanning distinct microarray platforms, we tested the hypothesis that nuclear-encoded FAO suppression represents a pan-subtype convergence point established at the earliest detectable disease stage. The complete analytical pipeline is summarised in Fig. 1.

**Figure 1.**
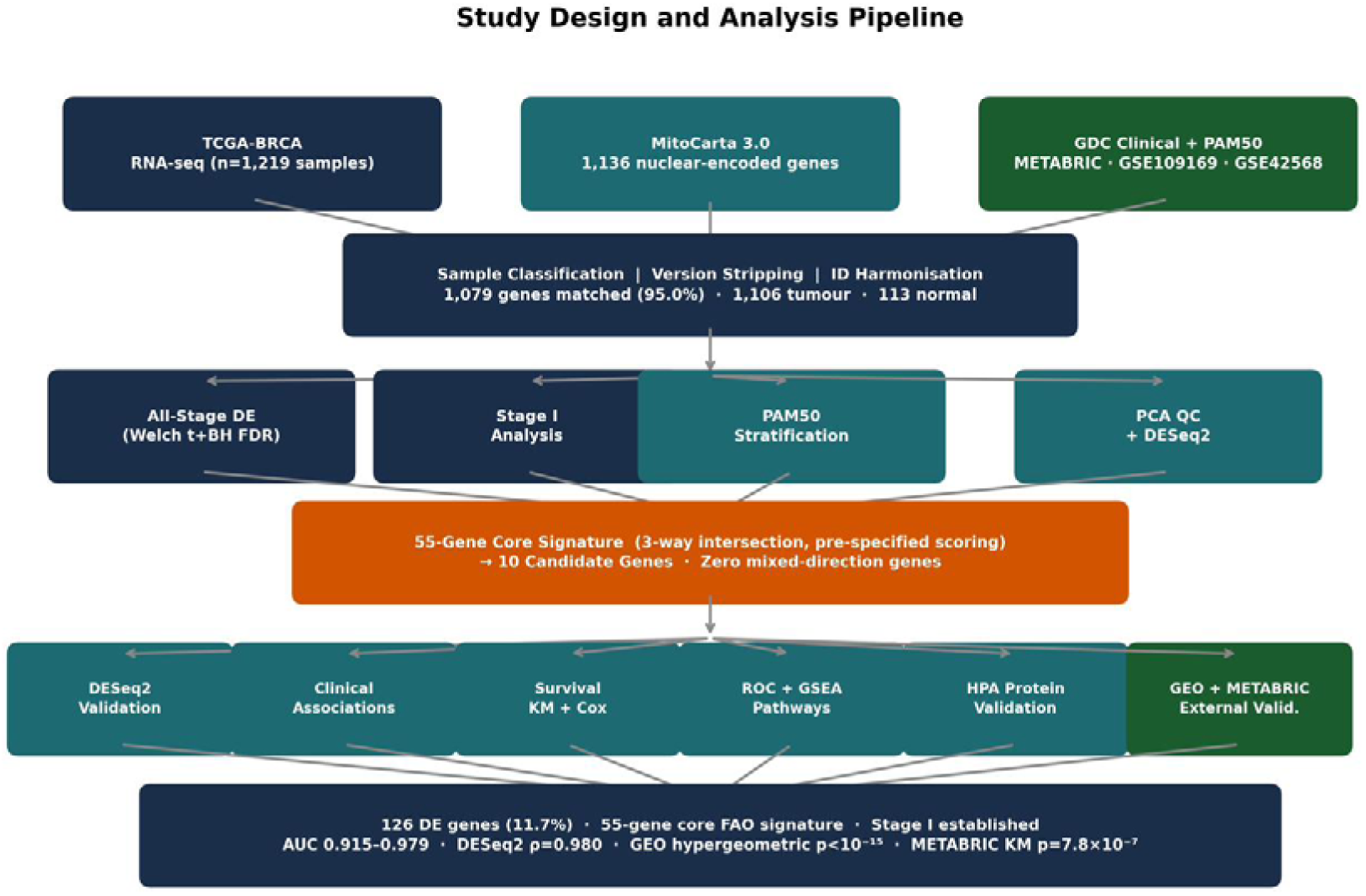
Study design and analysis pipeline. Schematic of the multi-stage analytical strategy. TCGA-BRCA RNA-seq data (n = 1,219 samples; GDC v09-07-2024) were intersected with 1,079 matched MitoCarta 3.0 nuclear-encoded mitochondrial genes, with PAM50 and clinical annotations from the TCGA Pan-Cancer Atlas. Following sample classification and principal component analysis (PCA) quality control, four complementary differential expression analyses yielded the 55-gene core fatty acid oxidation (FAO) signature. Pre-specified scoring identified 10 candidate genes prior to any clinical testing. Downstream characterisation included survival analysis, receiver operating characteristic (ROC) diagnostic performance, gene set enrichment analysis (GSEA) pathway enrichment, clinical association analysis, and external validation in two independent GEO cohorts (GSE109169, GSE42568). METABRIC was used for prognostic validation only. DE: differential expression; FDR: false discovery rate.

## Methods

### Data acquisition and preprocessing

RNA-seq gene expression data for TCGA-BRCA (GDC dataset version 09-07-2024; n = 1,226 samples; STAR-aligned, FPKM-UQ normalised) were downloaded via the GDC Data Portal [28]. Samples were classified by GDC sample code: primary tumour (code 01; n = 1,106), normal-adjacent tissue (code 11; n = 113), and metastatic (code 06; n = 7; excluded from all analyses). Nuclear-encoded mitochondrial genes were defined using MitoCarta 3.0 [10]. Ensembl gene IDs were harmonised by stripping version suffixes, yielding 1,079 matched genes (95.0% coverage). PAM50 subtype annotations were obtained from the TCGA Pan-Cancer Atlas phenotype file (n = 966 annotated tumours) [11]. Clinical data (AJCC stage, T stage, nodal status, ER status, HER2 status, age at diagnosis) were aligned to all 1,219 expression samples (100% alignment rate).

### Quality control

Principal component analysis (PCA) was performed on log₂-transformed FPKM-UQ+1 expression values across all 1,219 samples prior to any differential expression analysis, to confirm biological coherence of tumour versus normal groupings, identify potential outliers, and detect batch structure that could confound downstream analyses [12].

### Differential expression analysis

Differential expression was assessed using the Welch two-sample t-test applied to log₂(FPKM-UQ+1) expression values, with Benjamini–Hochberg (BH) false discovery rate (FDR) correction across all 1,079 tested genes [13, 14]. Significance thresholds: adjusted p < 0.05 and |log₂ fold change (FC)| > 1. Three complementary analyses were performed: (i) all-stage (1,106 tumours vs 113 normals); (ii) Stage I only (163 tumours vs 113 normals); and (iii) PAM50 subtype-stratified (LumA n = 427, LumB n = 192, Basal n = 144, HER2 n = 67, each vs 113 normals). Genes significant across all four PAM50 subtypes in concordant direction were designated pan-PAM50 differentially expressed.

### Core signature derivation

A 55-gene core signature was derived by three-way intersection of: (i) all-stage differentially expressed (DE) genes (n = 126); (ii) Stage I DE genes (n = 109); and (iii) pan-PAM50 DE genes (n = 57). Intersection required concordant directional change. A heatmap was generated for the top 50 core genes, with expression z-scored relative to the normal tissue mean and standard deviation per gene.

### Candidate gene selection

Ten candidate genes were selected from the 55-gene core signature by pre-specified objective scoring applied prior to any clinical testing, to prevent selection–outcome circularity [15]:

*Score = 0.40 × |log₂FC (all-stage)| + 0.40 × min(|log₂FC| across PAM50 subtypes) + 0.20 × (1 − cross-subtype SD of log₂FC)*

Weights prioritised dysregulation magnitude (0.40), minimum subtype-specific fold change as a pan-subtype robustness measure (0.40), and cross-subtype expression consistency (0.20).

### Pathway enrichment analysis

GSEA was performed using the pre-ranked Welch t-statistic gene list and MitoCarta 3.0 pathway annotations [16]. Significance threshold: FDR < 0.25 per established convention [16]. GSEA was performed independently in GSE42568 to assess pathway-level replication.

### Composite FAO suppression score

The composite FAO suppression score was computed as the mean z-score across the 10 candidate genes (relative to normal tissue mean and standard deviation), sign-inverted such that higher scores indicate greater FAO suppression. Derived exclusively in TCGA-BRCA and applied without modification to all validation cohorts.

### Clinical association analysis

Association of each candidate gene’s expression (median-dichotomised within tumours) with clinical variables (PAM50, AJCC stage, T stage, nodal status, ER status, HER2 status, age) was assessed using Chi-square tests for categorical variables and Mann-Whitney U for age. BH FDR correction was applied across all 70 simultaneous tests [13].

### Survival analysis

This prognostic biomarker study was conducted and reported in accordance with the REMARK guidelines [30]. Kaplan–Meier (KM) overall survival analysis was performed for each candidate gene individually (TCGA-BRCA; median-dichotomised; log-rank test) and for the composite FAO suppression score (METABRIC; n = 1,980; log-rank test) [17, 18]. Multivariable Cox proportional hazards regression was performed in METABRIC, adjusting for age, PAM50 subtype, and ER status [19]. The REMARK checklist is provided as Additional file 2.

### Diagnostic performance

Receiver operating characteristic (ROC) curves and area under the curve (AUC) were computed for each candidate gene against the binary tumour/normal classification in TCGA-BRCA using standard trapezoidal integration [20].

### Mutation correlation analysis

Expression of each candidate gene was compared between tumours with and without PIK3CA or TP53 somatic mutations (TCGA whole-exome sequencing data) using Welch t-test with BH FDR correction across 20 simultaneous tests.

### External validation cohorts

Two independent GEO cohorts [29] were used: GSE42568 (Affymetrix HG-U133 Plus 2.0, GPL570; 104 primary tumours, 17 normal-adjacent tissues) as primary replication; and GSE109169 (Affymetrix HuEx-1.0-ST, GPL5175; 25 matched tumour-normal pairs) as cross-platform validation. Core gene replication was assessed by hypergeometric test; effect size concordance by Spearman rank correlation of log₂FC vectors. The composite FAO score was applied without modification. METABRIC (n = 1,980; Illumina HT-12 v3; 1,143 OS events; median follow-up ∼10 years) [18], obtained from cBioPortal [27], was reserved for prognostic validation only, not DE replication, to prevent circularity with PAM50-based gene selection criteria.

### Methodological cross-validation

DESeq2 [21] was applied to raw TCGA-BRCA STAR count data as an independent cross-validation of the primary Welch t-test analysis. Duplicate gene IDs were resolved by retaining the entry with the highest total count. Spearman rank correlation of log₂FC estimates was computed across n = 1,074 testable genes.

### Statistical environment and AI disclosure

All analyses were implemented in Python 3.10 (Google Colab). Key packages: pandas 2.x, numpy 1.x, scipy.stats, scikit-learn, statsmodels, matplotlib, seaborn, pydeseq2. Analysis code is available at: https://github.com/y7ktdd4cjd-lgtm/breast-cancer-mitochondrial-FAO

AI-assisted tools (Claude, Anthropic) were used for manuscript drafting and code editing assistance. All analytical decisions, scientific interpretations, and final manuscript content were reviewed, verified, and approved by the authors. Use of AI assistance is disclosed in accordance with Springer Nature policy.

## Results

### Dataset composition and quality control

After exclusion of 7 metastatic samples, the discovery cohort comprised 1,106 primary breast tumours and 113 normal-adjacent tissues (Table 1). Of 1,136 MitoCarta 3.0 genes, 1,079 (95.0%) were matched following Ensembl ID harmonisation. PCA demonstrated clear separation between tumour and normal samples along PC1 (29.9% variance), with PC2 (10.6% variance) capturing inter-tumour heterogeneity consistent with subtype-level variation (Fig. 2a). No major batch structure or outlier clusters were identified.

**Figure 2.**
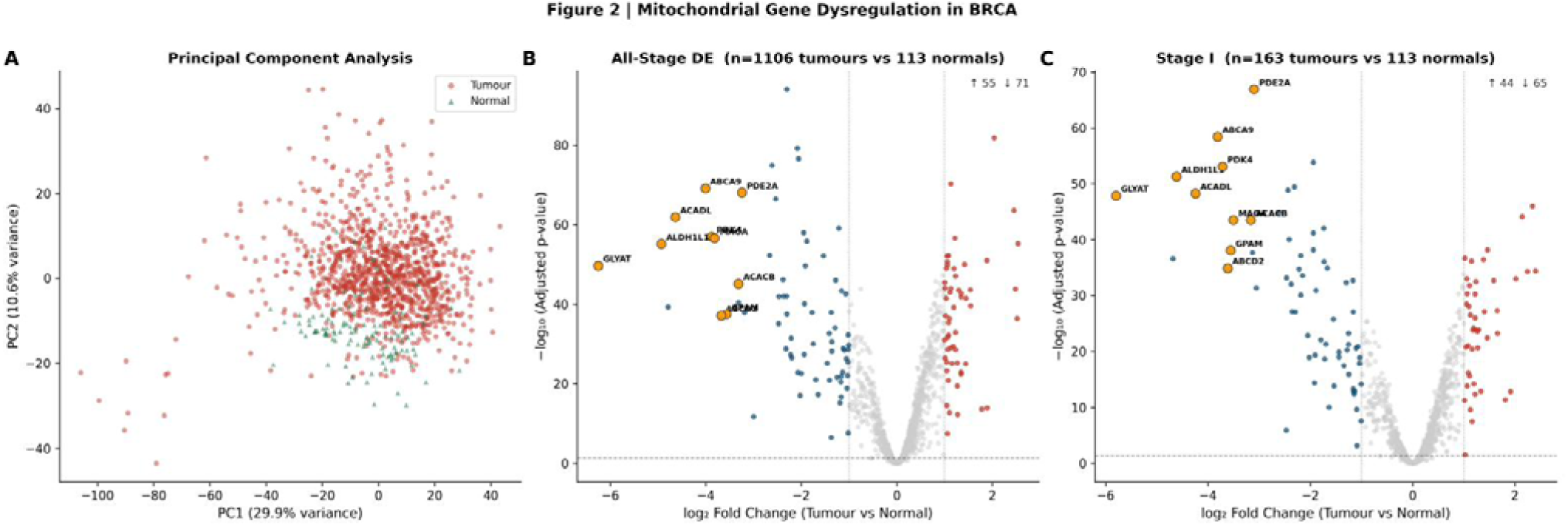
Mitochondrial gene dysregulation in breast cancer. (a) PCA of 1,219 TCGA-BRCA samples across 1,079 nuclear-encoded mitochondrial genes. Tumour samples (pink circles) and normal-adjacent tissues (teal triangles) separate along PC1 (29.9% variance explained). (b) Volcano plot of all-stage differential expression (1,106 tumours vs 113 normals; Welch t-test; BH FDR). Orange circles indicate the 10 candidate genes. Dotted lines: |log₂FC| = 1 and FDR = 0.05 thresholds. (c) Stage I-restricted volcano plot (163 Stage I tumours vs 113 normals), annotated as in (b). BH: Benjamini–Hochberg; FAO: fatty acid oxidation; FC: fold change; FDR: false discovery rate; PCA: principal component analysis.

**Table 1.**
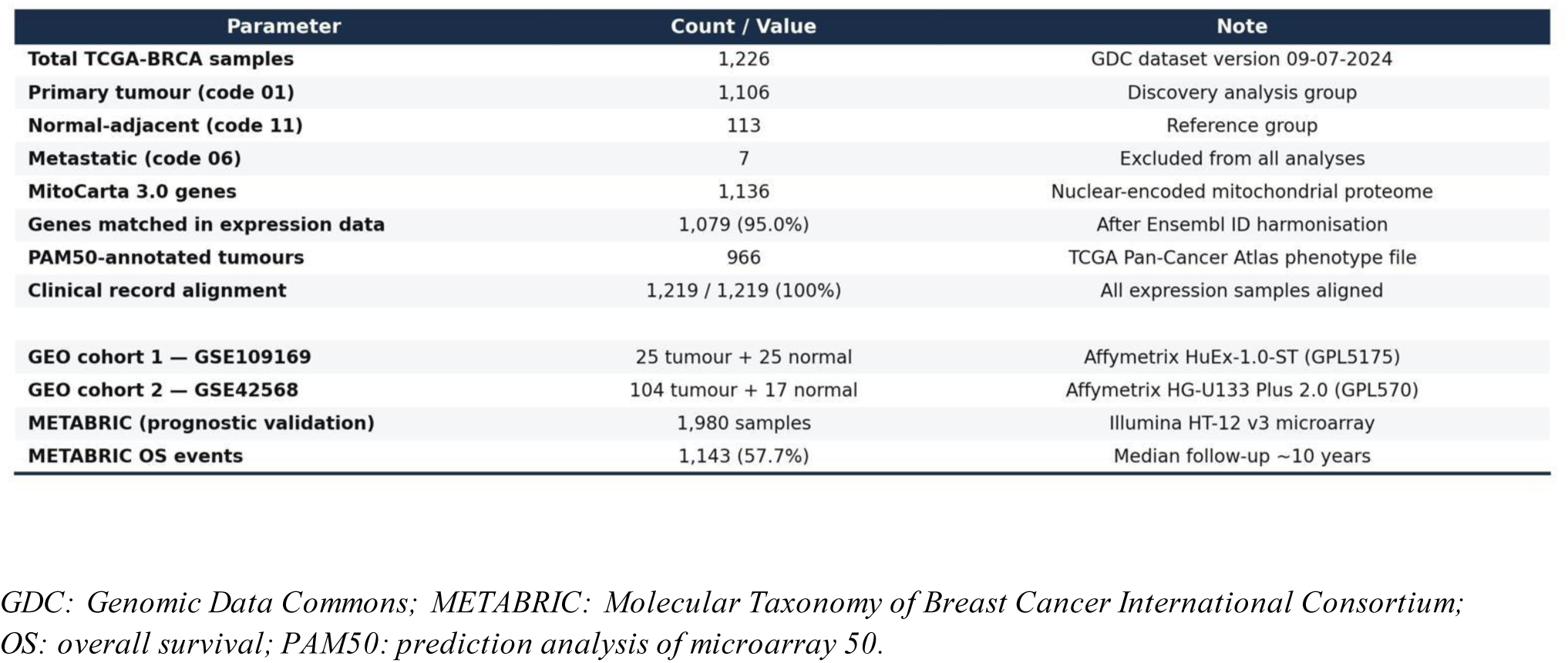
TCGA-BRCA dataset composition and validation cohort summary.

### Pervasive downregulation of nuclear-encoded mitochondrial genes across all stages

All-stage differential expression analysis identified 126 significantly dysregulated genes (11.7%): 55 upregulated and 71 downregulated in tumour (Fig. 2b). The predominance of downregulated genes indicates a net suppression of mitochondrial metabolic capacity. Stage I-restricted analysis identified 109 differentially expressed genes (44 upregulated, 65 downregulated; Fig. 2c), confirming that the majority of nuclear-encoded mitochondrial dysregulation is detectable at the earliest disease stage.

The near-complete recapitulation of all-stage differential expression in Stage I tumours — before lymph node involvement, metastatic spread, or therapeutic intervention — confirms that nuclear-encoded mitochondrial dysregulation is an early and consistent feature of the malignant state. This temporal coincidence of FAO suppression with the earliest detectable oncogenic stage is consistent with, though not sufficient to establish, a contributor role for metabolic reprogramming in breast cancer initiation. The uniformity of this signal across Stage I tumours of varying molecular backgrounds suggests that once a cell has undergone malignant transformation, FAO suppression is already present as a defining feature.

### Three-way intersection identifies a 55-gene pan-subtype core mitochondrial signature

PAM50-stratified analysis identified 57 genes consistently dysregulated across all four intrinsic subtypes in concordant direction (Additional file 1: Fig. S6). Three-way intersection yielded a 55-gene core signature (Fig. 3a). Critically, zero genes exhibited mixed directional change, indicating complete directional coherence. Heatmap visualisation of the top 50 core genes demonstrated uniform suppression across all PAM50 subtypes (Fig. 3b), with no subtype showing recovery toward normal expression levels.

**Figure 3.**
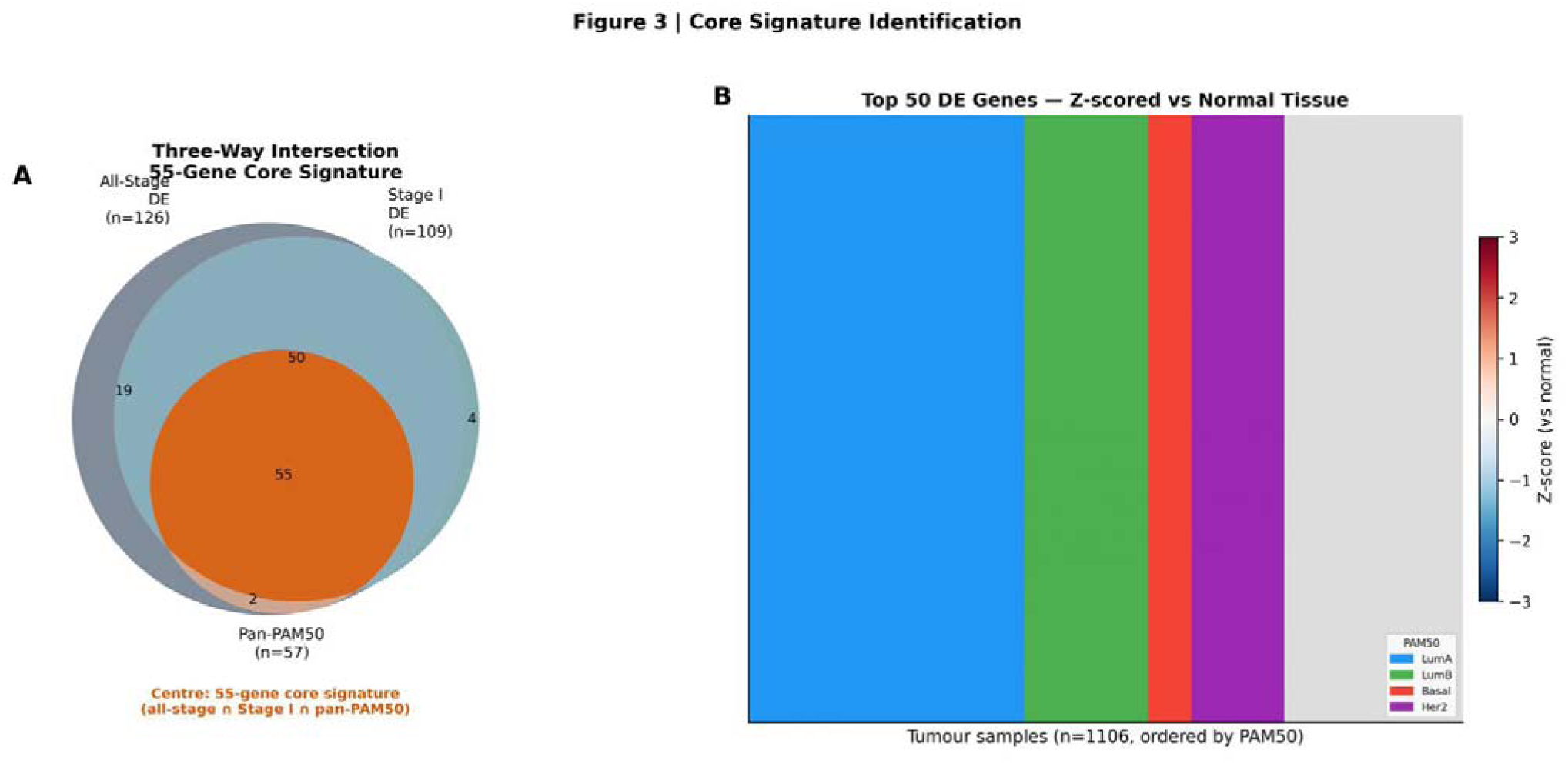
Core FAO signature identification. (a) Three-way Venn diagram of all-stage differentially expressed genes (n = 126), Stage I differentially expressed genes (n = 109), and pan-PAM50 differentially expressed genes (n = 57). The centre (orange) represents the 55-gene core FAO signature. (b) Heatmap of the top 50 differentially expressed core genes across all 1,106 tumour samples ordered by PAM50 subtype (LumA, LumB, Basal, HER2). Expression is z-scored relative to the normal tissue mean and standard deviation per gene. FAO: fatty acid oxidation; DE: differentially expressed.

### Fatty acid oxidation is the most significantly depleted mitochondrial pathway

GSEA identified fatty acid oxidation as the most significantly depleted mitochondrial pathway (NES = −2.130; FDR = 0.001), alongside xenobiotic metabolism (NES = −2.158; FDR = 0.001) and detoxification (NES = −1.884; FDR = 0.012) (Fig. 4a; Table 4). Fourteen pathways were enriched in tumour, including mitochondrial ribosome (NES = 2.073; FDR = 0.001), mitochondrial central dogma (NES = 2.013; FDR = 0.001), and OXPHOS assembly factors (NES = 1.553; FDR = 0.129). The composite FAO suppression score differed significantly across PAM50 subtypes (one-way ANOVA, p = 1.54 × 10⁻²²; Fig. 4b).

**Figure 4.**
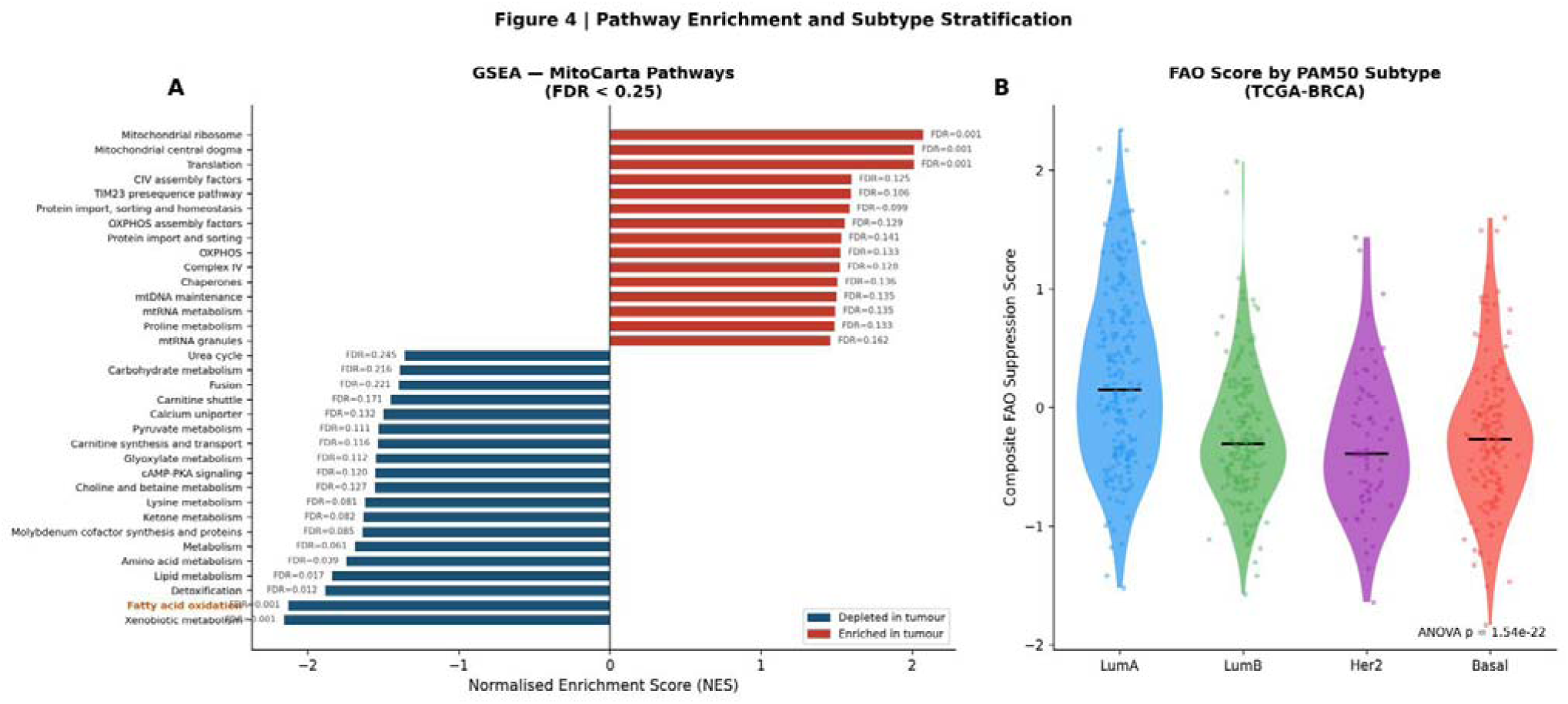
Pathway enrichment and FAO suppression score by subtype. (a) GSEA bar chart of MitoCarta 3.0 pathways significant at FDR < 0.25. Blue bars: depleted in tumour; red bars: enriched in tumour. Fatty acid oxidation (orange label) is the most significantly depleted metabolic pathway (NES = −2.130; FDR = 0.001). (b) Composite FAO suppression score by PAM50 subtype (TCGA-BRCA; n = 1,106 tumours; one-way ANOVA p = 1.54 × 10⁻²²). Horizontal bars indicate medians. FAO: fatty acid oxidation; FDR: false discovery rate; GSEA: gene set enrichment analysis; NES: normalised enrichment score; OXPHOS: oxidative phosphorylation.

The reciprocal enrichment pattern — suppression of FAO, lipid metabolism, and amino acid catabolism alongside upregulation of mitochondrial ribosome biogenesis, OXPHOS assembly, and mitochondrial RNA metabolism — indicates a selectively remodelled mitochondrion that has suppressed oxidative substrate catabolism while retaining and expanding its structural and translational machinery. This is consistent with the emerging understanding that tumour mitochondria remain bioenergetically active even as classical fuel oxidation pathways are downregulated [6, 22]. The pattern reflects a coordinated metabolic shift prioritising anabolic permissiveness over catabolic efficiency.

### Pre-specified candidate gene scoring selects 10 robust and functionally interpretable candidates

The pre-specified scoring formula ranked GLYAT first (score = 0.958; log₂FC = −6.25 all-stage; log₂FC = −5.80 Stage I), followed by ALDH1L1 (0.755), ACADL (0.671), ABCA9 (0.577), GPAM (0.569), PDK4 (0.553), ABCD2 (0.535), MAOA (0.523), PDE2A (0.490), and ACACB (0.488) (Table 2). The panel spans direct FAO enzymes (ACADL, ACACB), peroxisomal FAO transport (ABCD2), the pyruvate-FAO substrate gatekeeper PDK4, lipid transport mediators (ABCA9, GPAM), and FAO-linked redox enzymes (MAOA, ALDH1L1, GLYAT).

**Table 2.**
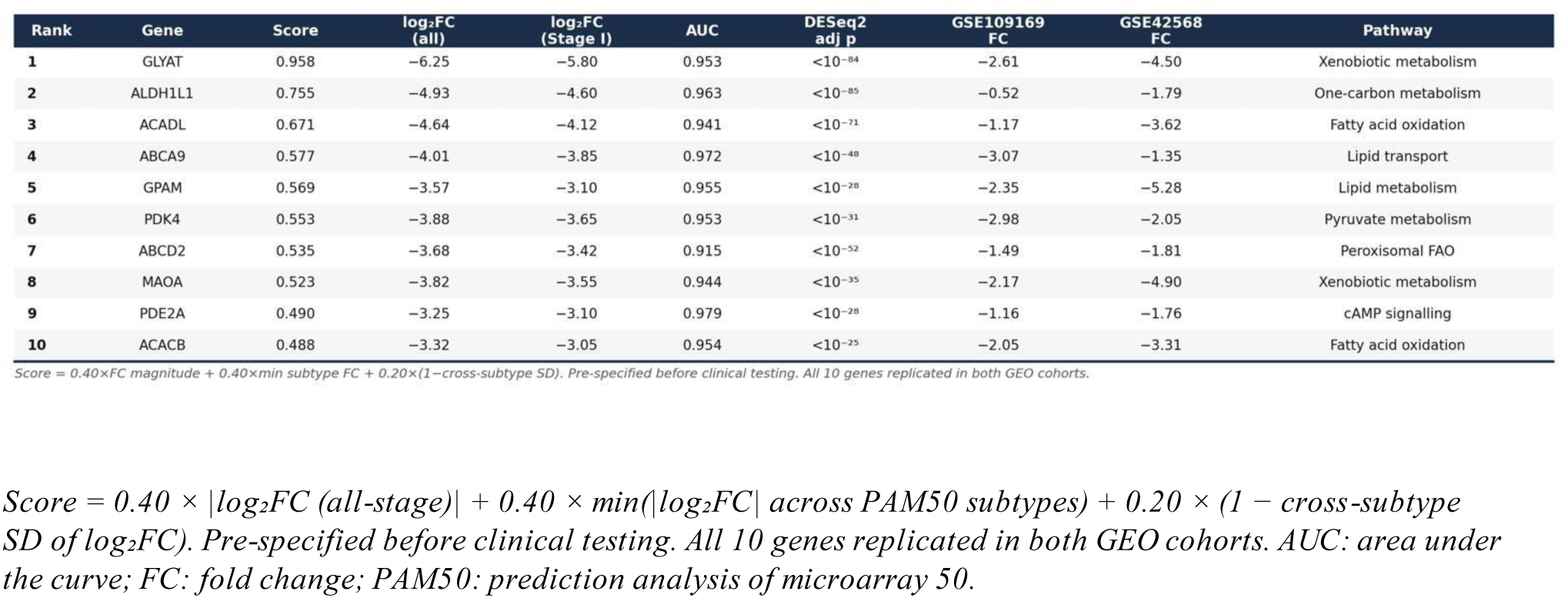
Top 10 candidate genes selected by pre-specified objective scoring.

### Candidate genes discriminate tumour from normal tissue with high diagnostic accuracy

ROC analysis yielded AUC values of 0.915 (ABCD2) to 0.979 (PDE2A), with all 10 genes exceeding 0.90 (Fig. 5a). The composite FAO suppression score achieved AUC = 0.989 (p = 1.19 × 10⁻¹⁰) in GSE42568 and AUC = 0.998 (p = 1.60 × 10⁻⁹) in GSE109169, demonstrating near-complete separation of tumour and normal tissue in independent cohorts.

**Figure 5.**
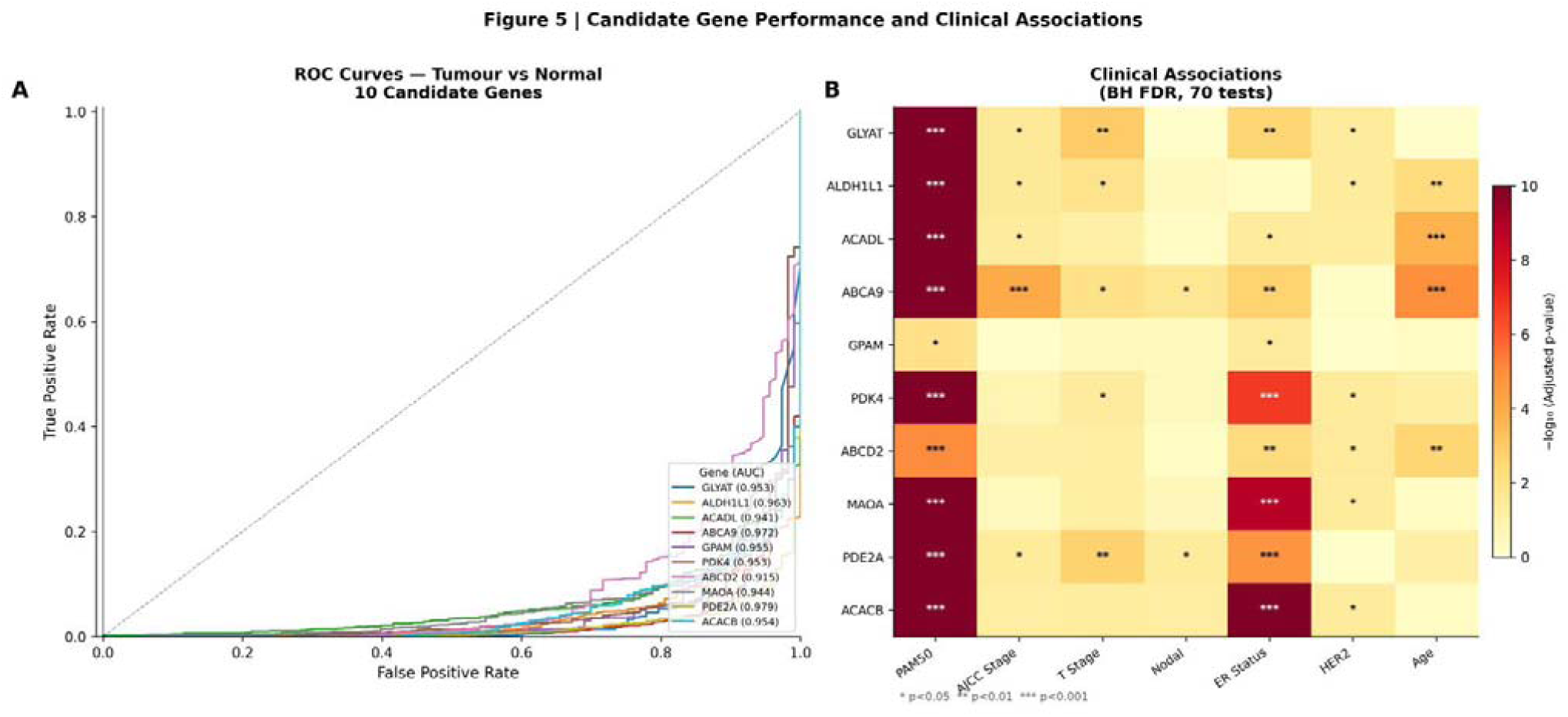
Candidate gene diagnostic performance and clinical associations. (a) ROC curves for the 10 candidate genes (tumour vs normal classification; TCGA-BRCA). AUC values are shown in the legend. (b) Clinical association heatmap across 10 candidate genes and 7 clinical variables. Colour intensity represents −log₁₀(adjusted p-value). Asterisks denote BH FDR significance thresholds after correction across 70 simultaneous tests: *adj p < 0.05; **adj p < 0.01; ***adj p < 0.001. AUC: area under the curve; BH: Benjamini–Hochberg; FDR: false discovery rate; ROC: receiver operating characteristic.

### Candidate genes are broadly associated with clinical and pathological features

Clinical association analysis across 70 simultaneous tests (BH FDR) identified significant PAM50 associations in all 10 candidate genes (all adjusted p < 0.001; Fig. 5b; Table 3). Association breadth ranged from 2/7 clinical variables (GPAM) to 6/7 (ABCA9). AJCC stage associations were significant for GLYAT, ALDH1L1, ABCA9, and PDE2A.

**Table 3.**
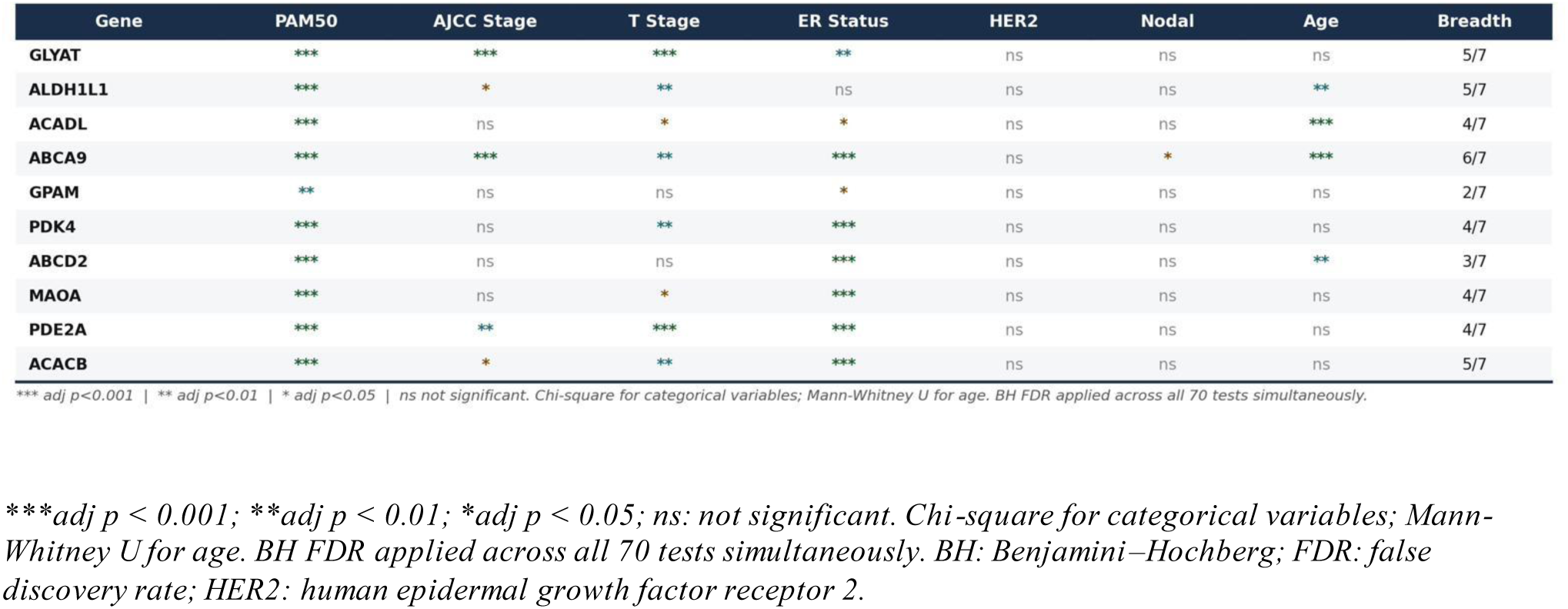
Clinical association analysis (70 tests, BH FDR correction).

### FAO suppression associates with overall survival

KM analysis in TCGA-BRCA identified significant overall survival associations for ACADL (log-rank p = 0.040) and ABCD2 (p = 0.028), with higher expression associated with superior survival (Fig. 6a; Additional file 1: Figs. S1, S2). In METABRIC, the composite FAO suppression score provided strong survival separation (log-rank p = 7.82 × 10⁻⁷; Fig. 6b). In multivariable Cox regression adjusting for age, PAM50, and ER status, MAOA achieved independent prognostic significance (HR = 0.890, 95% CI 0.819–0.967; adjusted p = 0.009), while the composite FAO score did not reach adjusted significance (HR = 0.942, 95% CI 0.881–1.007; p = 0.078; Fig. 6c).

**Figure 6.**
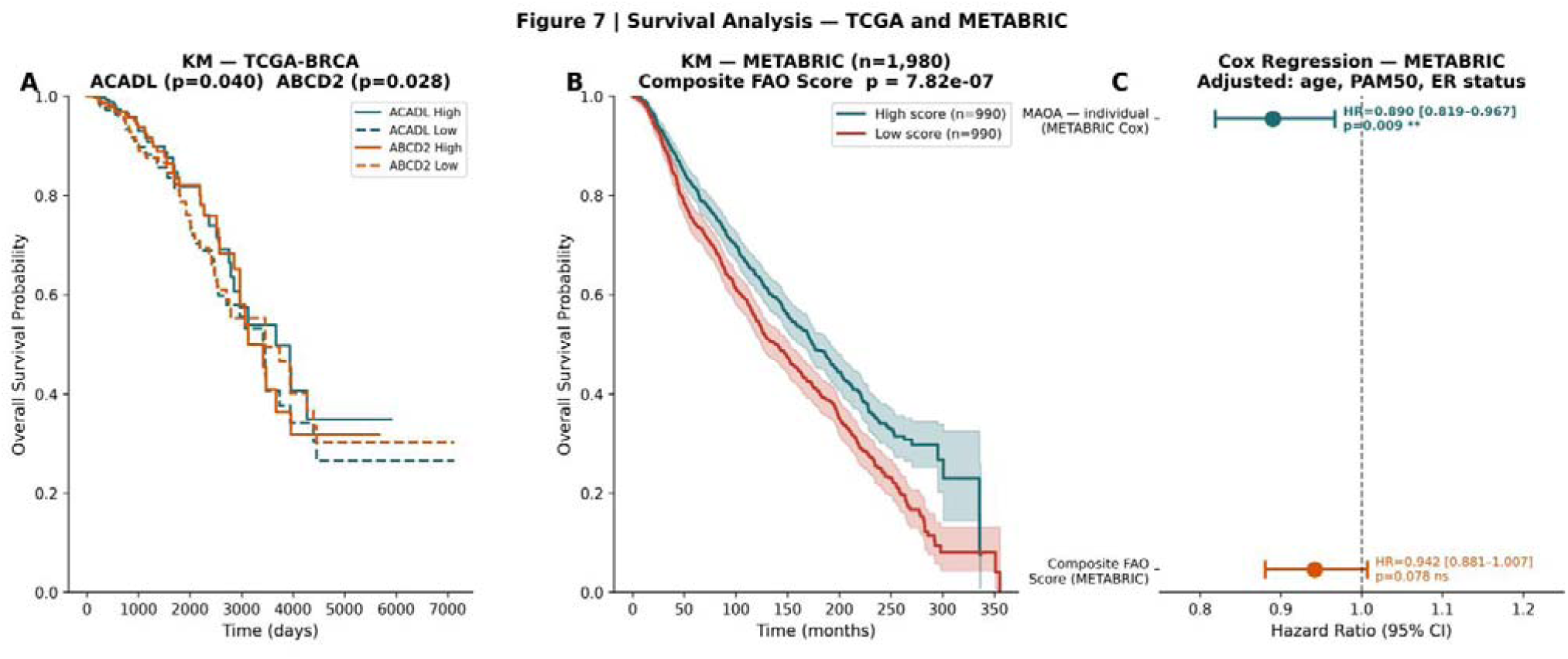
Survival analysis in TCGA-BRCA and METABRIC. (a) Kaplan–Meier overall survival curves for ACADL (log-rank p = 0.040) and ABCD2 (log-rank p = 0.028) in TCGA-BRCA (median-dichotomised expression). (b) Kaplan–Meier overall survival curves by composite FAO suppression score (high vs low; METABRIC; n = 1,980; log-rank p = 7.82 × 10⁻⁷). Shading indicates 95% confidence intervals. (c) Forest plot of multivariable Cox proportional hazards regression in METABRIC (adjusted: age, PAM50 subtype, ER status). MAOA HR = 0.890 (95% CI 0.819–0.967; adj p = 0.009); composite FAO suppression score HR = 0.942 (95% CI 0.881–1.007; p = 0.078). CI: confidence interval; ER: oestrogen receptor; FAO: fatty acid oxidation; HR: hazard ratio; KM: Kaplan–Meier.

The KM and Cox results address different questions and are not contradictory. The log-rank test assesses unadjusted survival separation, while Cox regression partitions prognostic variance among covariates. Both PAM50 and ER status are strong prognostic variables correlated with FAO suppression score [2], so attenuation after adjustment is expected. A HR of 0.942 (95% CI 0.881–1.007) should be interpreted as a directionally consistent trend rather than a negative result. The independent significance of MAOA may reflect a prognostic contribution specific to this gene not fully captured by subtype classification. All survival findings are best characterised as hypothesis-generating, consistent with REMARK guidelines [30].

### FAO suppression is largely independent of PIK3CA but modulated by TP53 mutation status

Expression of the 10 candidate genes was largely independent of PIK3CA mutation status (2/10 genes significant at BH FDR < 0.05: ABCA9 and GPAM; Fig. 7a). In contrast, 9/10 candidate genes showed significantly altered expression between TP53-mutated and TP53-wildtype tumours (Fig. 7b), with TP53-mutant tumours exhibiting deeper FAO suppression.

**Figure 7.**
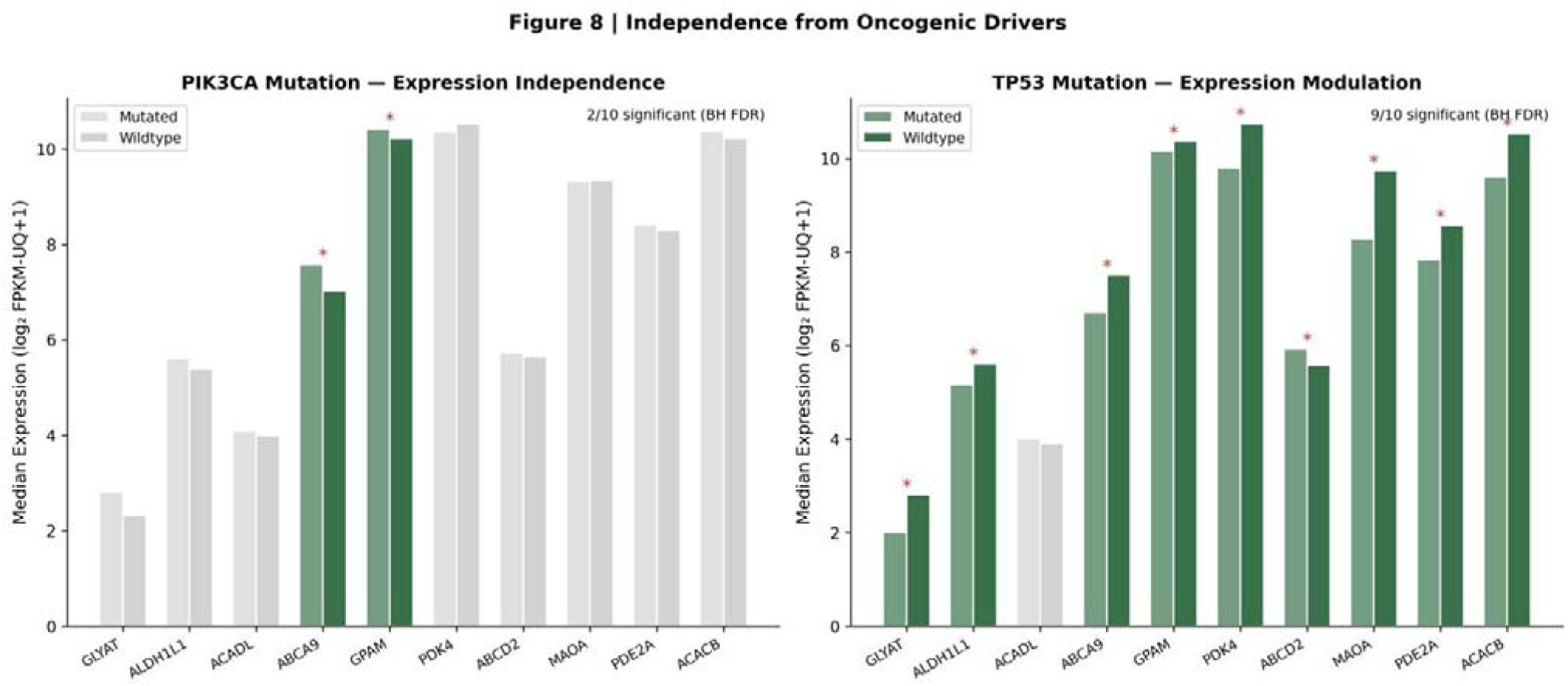
Candidate gene expression independence from oncogenic drivers. Median expression (log₂ FPKM-UQ+1) of 10 candidate genes in TCGA-BRCA tumours stratified by (a) PIK3CA mutation status (mutated vs wildtype; 2/10 genes significant at BH FDR < 0.05; red asterisks) and (b) TP53 mutation status (9/10 genes significant). Dark bars: mutated; light bars: wildtype. BH: Benjamini–Hochberg; FDR: false discovery rate; FPKM-UQ: fragments per kilobase per million reads, upper quartile normalised.

The association of 9/10 candidate genes with TP53 mutation status reflects the well-established role of wild-type p53 as a transcriptional activator of FAO-related genes including LPIN1 and CPT1C [23]. Loss of this regulatory function upon TP53 mutation would be expected to deepen, rather than initiate, suppression: candidate gene suppression is present across all tumours regardless of TP53 status, and is accentuated in its magnitude in the context of TP53 loss. TP53 mutation may therefore represent one of several upstream mechanisms that converge on FAO suppression as a shared metabolic endpoint.

### Cross-platform external validation confirms robustness of the core signature

In GSE42568 (104 tumours, 17 normals), 34/52 testable core genes were replicated (65%; expected 8.6 by chance; fold enrichment 3.9×; hypergeometric p = 4.88 × 10⁻¹⁶; 100% directional concordance; Fig. 8b; Additional file 1: Fig. S4). All 10 candidate genes individually replicated. The Spearman correlation of cross-cohort effect sizes was ρ = 0.542 (p = 3.34 × 10⁻⁵; Fig. 8a). Composite AUC = 0.989. GSEA confirmed FAO as a significantly depleted pathway in GSE42568 (NES = −2.064; FDR = 0.001; Table 4), with 14/34 MitoCarta pathways replicated directionally.

**Figure 8.**
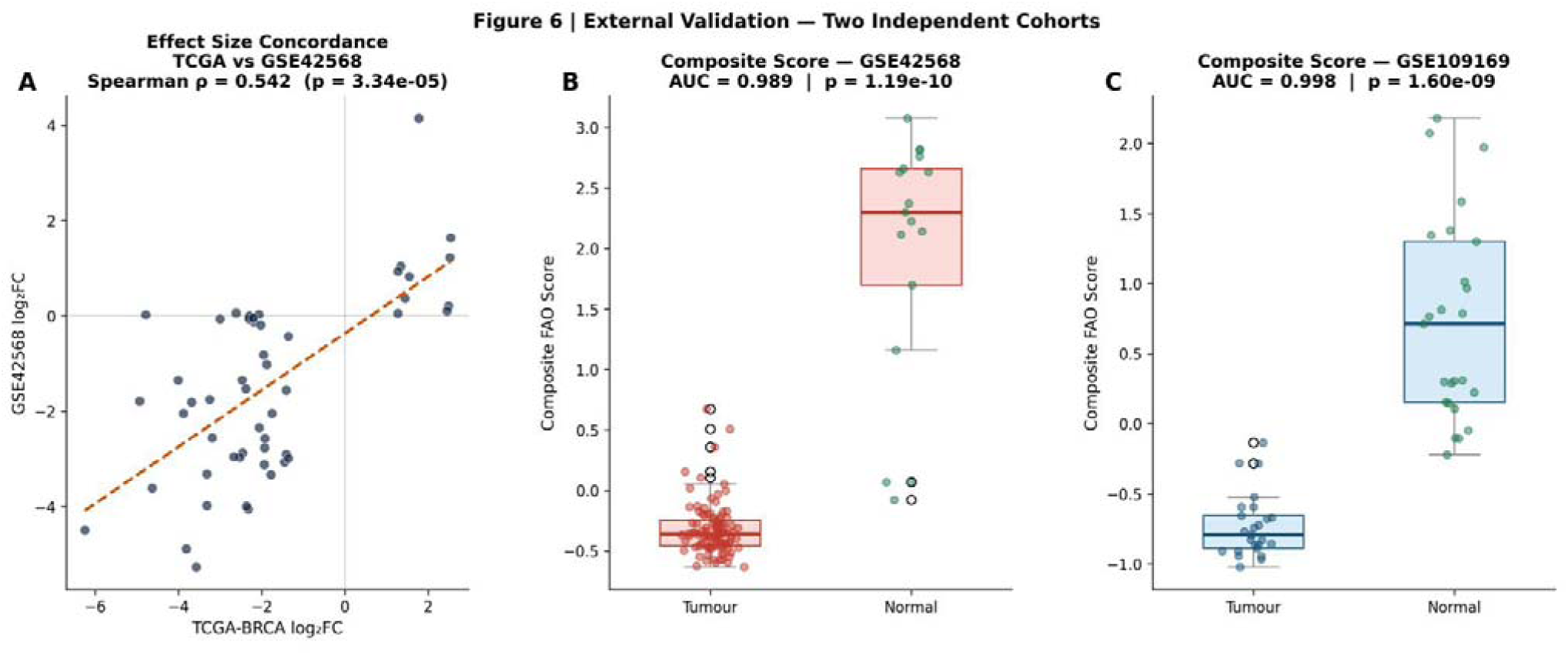
External validation in two independent cohorts. (a) Spearman rank correlation of log₂ fold-change effect sizes between TCGA-BRCA and GSE42568 across 52 testable core genes (ρ = 0.542; p = 3.34 × 10⁻⁵). Dashed line: Deming regression. (b) Composite FAO suppression score in GSE42568 (104 tumours vs 17 normals; AUC = 0.989; p = 1.19 × 10⁻¹⁰). Open circles indicate outliers. (c) Composite FAO suppression score in GSE109169 (25 tumours vs 25 normals; cross-platform; AUC = 0.998; p = 1.60 × 10⁻⁹). AUC: area under the curve; FAO: fatty acid oxidation.

**Table 4.** GSEA pathway enrichment — MitoCarta 3.0 pathways (FDR < 0.25).

| Pathway | NES | FDR | Direction | Replicated in GSE42568? | GSE42568 NES |
| --- | --- | --- | --- | --- | --- |
| Xenobiotic metabolism | -2.158 | 0.0010 | Depleted | ✓ Yes | -1.837 |
| Fatty acid oxidation | -2.130 | 0.0010 | Depleted | ✓ Yes | -2.064 |
| Detoxification | -1.884 | 0.0119 | Depleted | ✓ Yes | -1.875 |
| Lipid metabolism | -1.840 | 0.0171 | Depleted | ✓ Yes | -1.640 |
| Amino acid metabolism | -1.745 | 0.0392 | Depleted | ✓ Yes | -1.457 |
| Metabolism | -1.689 | 0.0613 | Depleted | ✓ Yes | -1.662 |
| Molybdenum cofactor synthesis and proteins | -1.640 | 0.0852 | Depleted | — | — |
| Ketone metabolism | -1.633 | 0.0820 | Depleted | ✓ Yes | -1.400 |
| Lysine metabolism | -1.621 | 0.0805 | Depleted | ✓ Yes | -1.443 |
| Choline and betaine metabolism | -1.558 | 0.1270 | Depleted | ✓ Yes | -1.477 |
| cAMP-PKA signaling | -1.554 | 0.1199 | Depleted | ✓ Yes | -0.748 |
| Glyoxylate metabolism | -1.551 | 0.1122 | Depleted | ✓ Yes | -1.165 |
| Carnitine synthesis and transport | -1.538 | 0.1156 | Depleted | ✓ Yes | -0.682 |
| Pyruvate metabolism | -1.533 | 0.1111 | Depleted | ✓ Yes | -1.056 |
| Calcium uniporter | -1.500 | 0.1324 | Depleted | ✓ Yes | -0.841 |
| Carnitine shuttle | -1.453 | 0.1705 | Depleted | — | — |
| Fusion | -1.398 | 0.2209 | Depleted | ✓ Yes | -0.984 |
| Carbohydrate metabolism | -1.393 | 0.2160 | Depleted | ✓ Yes | -1.512 |
| Urea cycle | -1.360 | 0.2446 | Depleted | ✓ Yes | 0.901 |
| mtRNA granules | 1.458 | 0.1620 | Enriched | ✓ Yes | 1.372 |
| Proline metabolism | 1.486 | 0.1326 | Enriched | — | — |
| mtRNA metabolism | 1.491 | 0.1351 | Enriched | ✓ Yes | 1.829 |
| mtDNA maintenance | 1.499 | 0.1346 | Enriched | ✓ Yes | 2.281 |
| Chaperones | 1.507 | 0.1362 | Enriched | ✓ Yes | 0.987 |
| Complex IV | 1.521 | 0.1280 | Enriched | ✓ Yes | -0.869 |
| OXPHOS | 1.526 | 0.1325 | Enriched | ✓ Yes | -1.547 |
| Protein import and sorting | 1.531 | 0.1412 | Enriched | ✓ Yes | 0.615 |
| OXPHOS assembly factors | 1.553 | 0.1289 | Enriched | ✓ Yes | -0.882 |
| Protein import, sorting and homeostasis | 1.586 | 0.0988 | Enriched | ✓ Yes | 0.822 |
| TIM23 presequence pathway | 1.595 | 0.1058 | Enriched | ✓ Yes | 0.530 |
| CIV assembly factors | 1.600 | 0.1255 | Enriched | ✓ Yes | 0.990 |
| Translation | 2.011 | 0.0010 | Enriched | ✓ Yes | 0.934 |
| Mitochondrial central dogma | 2.013 | 0.0010 | Enriched | ✓ Yes | 1.658 |
| Mitochondrial ribosome | 2.073 | 0.0010 | Enriched | ✓ Yes | -0.704 |

NES = normalised enrichment score. FDR<0.25 per Subramanian et al. 2005. Replicated = significant in independent GSE42568 GSEA.
NES: normalised enrichment score. $FDR \leq 0.25$ per Subramanian et al. [16]. Replicated = significant in independent GSE42568 GSEA. FAO: fatty acid oxidation; FDR: false discovery rate; GSEA: gene set enrichment analysis; OXPHOS: oxidative phosphorylation.

In GSE109169 (25 matched pairs; cross-platform), 19/43 testable core genes were replicated (44%; expected 1.3; fold enrichment 14.1×; hypergeometric p = 5.63 × 10⁻²¹; 100% directional concordance; Additional file 1: Fig. S3). All 10 candidate genes individually replicated. Composite AUC = 0.998 (Fig. 8c). The lower absolute replication rate in GSE109169 relative to GSE42568 reflects cross-platform constraints and reduced statistical power; fold enrichment (14.1×) and directional concordance (100%) confirm the signal is platform-independent (Table 5).

**Table 5.**
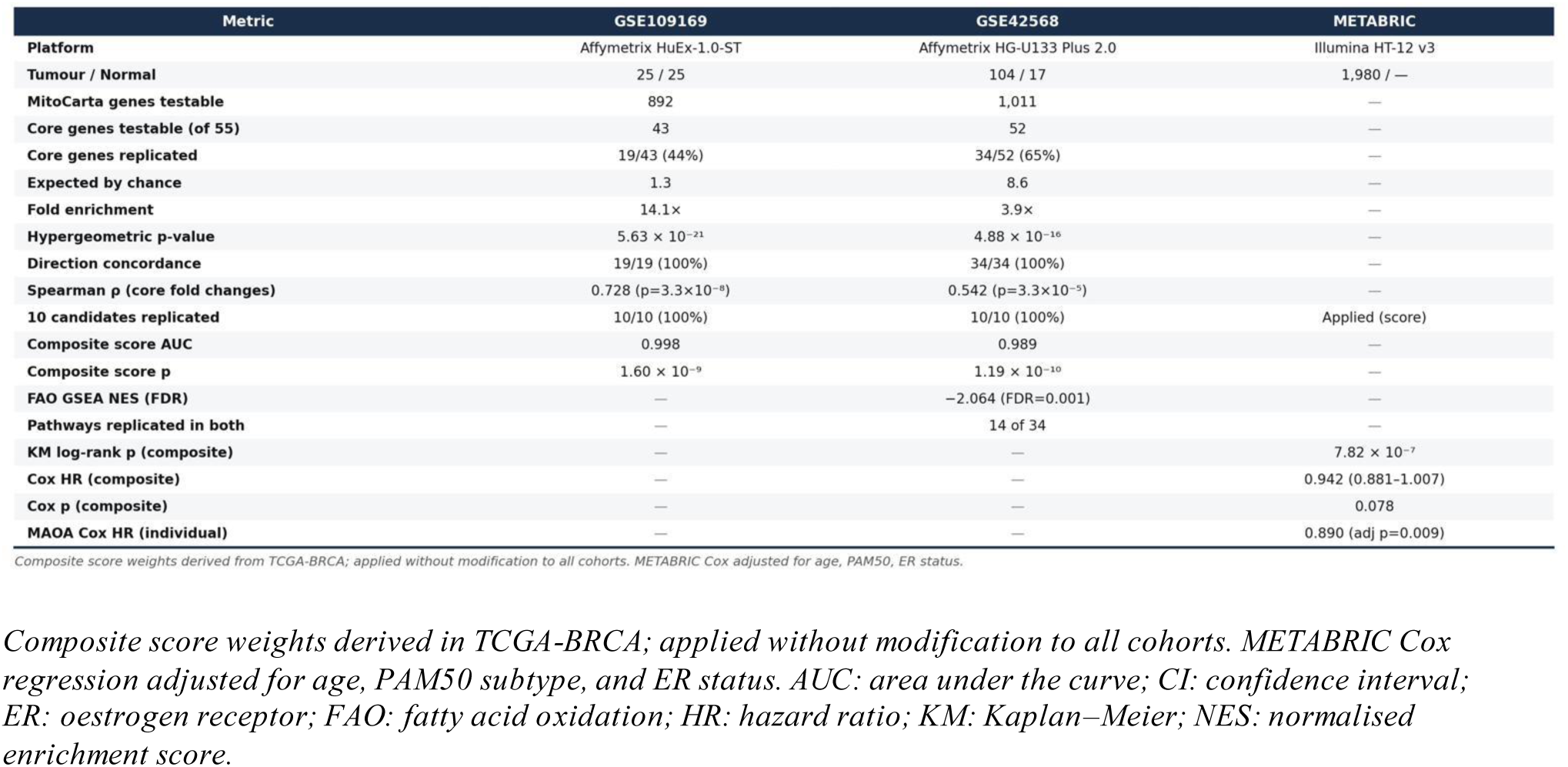
Cross-platform external validation summary.

### DESeq2 cross-validation confirms methodological robustness

DESeq2 [21] applied to raw TCGA-BRCA STAR count data produced Spearman ρ = 0.980 (p ≈ 0; n = 1,074 genes) between DESeq2 and Welch t-test log₂FC estimates (Additional file 1: Fig. S5). DESeq2 identified 187 significant genes, confirming all 126 Welch-significant genes. This concordance validates the primary Welch t-test analysis as methodologically robust.

## Discussion

The identification of a 10-gene nuclear-encoded mitochondrial signature — consistently, significantly, and uniformly suppressed across all molecular subtypes and detectable at Stage I — adds a metabolic dimension to our understanding of breast cancer initiation that complements the subtype-defining genetic events currently used to stratify treatment. That a convergent metabolic phenotype accompanies, and may precede, the full declaration of subtype-specific programmes suggests that malignant transformation requires not only nuclear permissiveness but a concurrent mitochondrial context. This observation does not challenge the somatic mutation model but extends it: FAO suppression may represent a co-occurring and potentially necessary metabolic condition for tumour establishment, one that is targetable independently of subtype-specific genomic features.

The finding that nuclear-encoded FAO is the most significantly depleted mitochondrial pathway, detectable at Stage I, and exhibiting 100% directional concordance across two independent platforms, is consistent with a model in which FAO suppression is an early and conserved metabolic reorganisation. The three-way intersection strategy — enforcing concordance across all-stage, Stage I, and all four PAM50 subtypes simultaneously — was specifically designed to recover this conserved signal while filtering subtype-specific and stage-dependent effects. The absence of any mixed-direction genes in the 55-gene core signature provides the most systematic transcriptomic evidence to date for pan-subtype FAO suppression at genome-wide scale in breast cancer.

Prior work has characterised subtype-specific alterations in fatty acid metabolism in breast cancer, with different intrinsic subtypes exhibiting distinct patterns of FAO enzyme expression [9]. The present study extends these observations by demonstrating that, beneath subtype-level variability, a convergent FAO suppression signature is detectable across all intrinsic subtypes simultaneously. Several converging mechanisms may explain why FAO suppression constitutes a shared endpoint. Cyclin D1, which is overexpressed across breast cancer subtypes as a pro-proliferative driver, directly represses PPARα transcriptional activity in breast cancer cells, inhibiting FAO gene expression [24]. Oncogene-driven de novo lipogenesis elevates malonyl-CoA, allosterically inhibiting CPT1 and preventing long-chain fatty acid import into the mitochondrion for β-oxidation [8]. Additionally, TP53 loss removes a key transcriptional activator of FAO-related genes including LPIN1 and CPT1C [23]. These mechanisms are not mutually exclusive and may operate redundantly to produce the convergent suppression observed.

The association of FAO suppression with TP53 mutation status across 9/10 candidate genes merits attention. TP53 is among the most frequently mutated tumour suppressor genes in breast cancer [25] and is nearly ubiquitously mutated in Basal-like tumours [25]. Wild-type p53 transcriptionally activates FAO-related genes, promoting mitochondrial FAO as a stress-adaptive programme [23]. Loss of this activation upon TP53 mutation would deepen FAO suppression. The pan-subtype character of the signature — extending to LumA tumours with relatively low TP53 mutation rates [25] — indicates that TP53 loss is a modulator of degree rather than a requisite initiating event.

The survival data require careful interpretation. The composite FAO suppression score separates METABRIC survival curves strongly (p = 7.82 × 10⁻⁷) but does not reach adjusted significance in multivariable Cox regression (p = 0.078), while MAOA achieves independent significance after full adjustment (HR = 0.890; adj p = 0.009). PAM50 and ER status are themselves strong prognostic variables correlated with FAO suppression score [2], so attenuation after adjustment is expected. The independent prognostic significance of MAOA may reflect a contribution not fully captured by subtype classification. Consistent with REMARK guidelines [30], all survival findings are best characterised as hypothesis-generating, warranting prospective evaluation in appropriately powered cohorts.

Several limitations warrant acknowledgement. First, TCGA-BRCA normal-adjacent tissue is derived from cancer-bearing individuals; field cancerisation effects may mean that differential expression magnitudes are conservative [26]. Second, transcriptomic data cannot account for post-transcriptional regulation, protein abundance, or enzymatic activity; whether FAO gene suppression translates to reduced FAO flux requires functional validation in primary tumour models. Third, the observational design precludes causal inference: the data establish co-occurrence of FAO suppression with malignant transformation at Stage I, but cannot determine whether suppression precedes and enables oncogenesis or is a parallel consequence. Fourth, the composite FAO score’s Cox regression result (p = 0.078) should be regarded as hypothesis-generating rather than evidence of validated prognostic utility. Finally, GSE109169 (25 matched pairs) is insufficiently powered for individual gene-level replication and contributes cross-platform directional evidence only.

## Conclusions

This study provides systematic, multi-cohort transcriptomic evidence that nuclear-encoded mitochondrial fatty acid oxidation suppression is a convergent, pan-subtype feature of breast cancer, detectable at Stage I and reproducible across independent platforms. The 55-gene core mitochondrial signature and the 10 pre-specified candidate genes collectively define a shared metabolic phenotype beneath the heterogeneity of breast cancer subtypes. Association with TP53 mutation status and cyclin D1-driven PPARα suppression identifies candidate mechanistic axes warranting experimental investigation. Further proteomic and functional validation is required before causality can be established; however, the present findings motivate a broader conceptual framework for breast cancer initiation — one that extends beyond subtype-specific genetic events to include convergent mitochondrial metabolic reprogramming. Biological phenomena are rarely reducible to a single mechanism, and the convergent mitochondrial signature identified here suggests that the origins of malignancy are no exception.

## Abbreviations

AUC: Area under the curve
BH: Benjamini–Hochberg
CI: Confidence interval
DE: Differentially expressed
ER: Oestrogen receptor
FAO: Fatty acid oxidation
FC: Fold change
FDR: False discovery rate
FPKM-UQ: Fragments per kilobase per million reads, upper quartile normalised
GDC: Genomic Data Commons
GSEA: Gene set enrichment analysis
HER2: Human epidermal growth factor receptor 2
HR: Hazard ratio
KM: Kaplan–Meier
NES: Normalised enrichment score
OS: Overall survival
OXPHOS: Oxidative phosphorylation
PAM50: Prediction analysis of microarray 50
PCA: Principal component analysis
ROC: Receiver operating characteristic
SD: Standard deviation
TCGA-BRCA: The Cancer Genome Atlas Breast Cancer.

## Declarations

### Ethics approval and consent to participate

This study used publicly available, de-identified data from TCGA [28], GEO [29], and METABRIC [18, 27] repositories. No ethical approval or participant consent was required.

### Consent for publication

Not applicable.

### Availability of data and materials

TCGA-BRCA data are publicly available via the GDC Data Portal (version 09-07-2024) [28]. GEO datasets GSE42568 and GSE109169 are available via the NCBI Gene Expression Omnibus [29]. METABRIC data were obtained from cBioPortal [27]. Analysis code is available at https://github.com/y7ktdd4cjd-lgtm/breast-cancer-mitochondrial-FAO

### Competing interests

The authors declare that they have no competing interests.

### Funding

Not applicable.

### Authors’ contributions

AG: conceptualisation, methodology, software, formal analysis, writing — original draft. HM: formal analysis, writing — review and editing. LM: investigation, writing — review and editing. All authors read and approved the final manuscript.

## Acknowledgements

The authors acknowledge the contributions of the TCGA Research Network, the NCBI Gene Expression Omnibus, and the METABRIC consortium in making these datasets publicly available.

## Additional Files

Additional file 2: REMARK checklist.

**Additional file 1: Figure S1.**
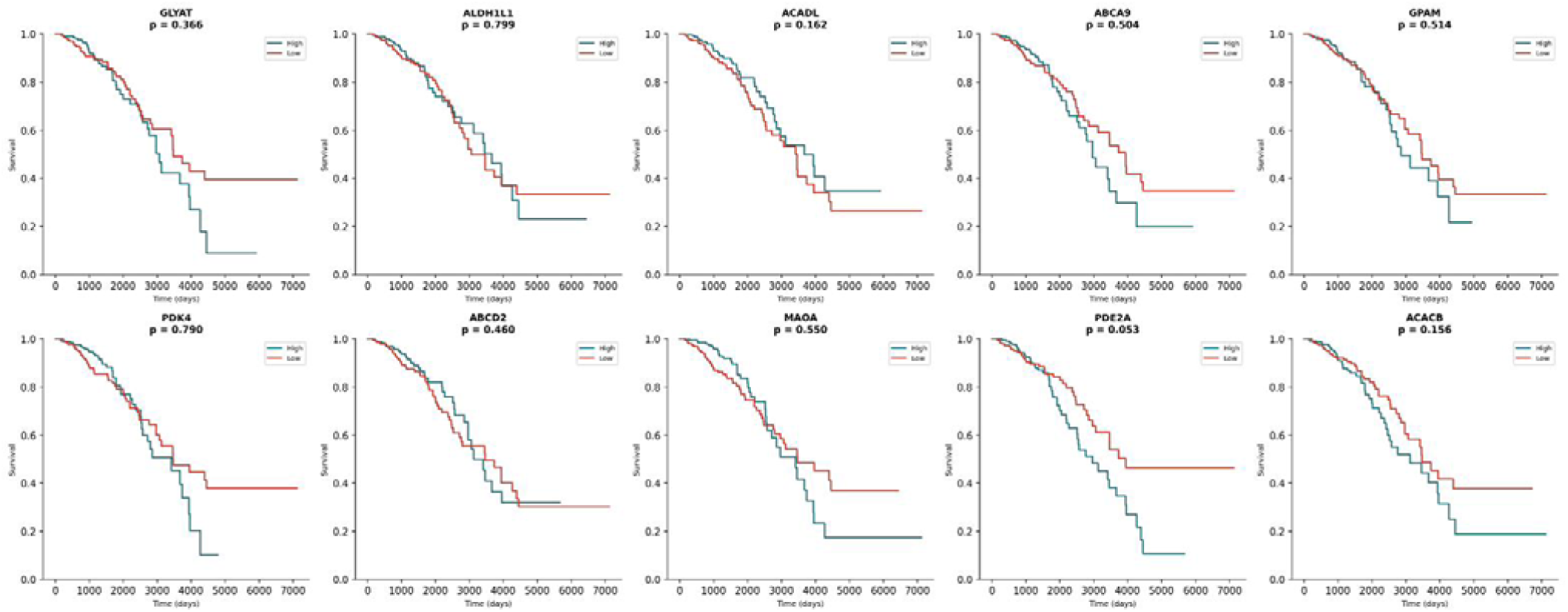
Kaplan–Meier overall survival — all 10 candidate genes (TCGA-BRCA). Kaplan–Meier overall survival curves for each of the 10 candidate genes individually (median-dichotomised expression within tumours; TCGA-BRCA; log-rank p-values shown). ACADL (p = 0.040) and ABCD2 (p = 0.028) reached significance. High expression: teal; low expression: red.

**Additional file 1: Figure S2.**
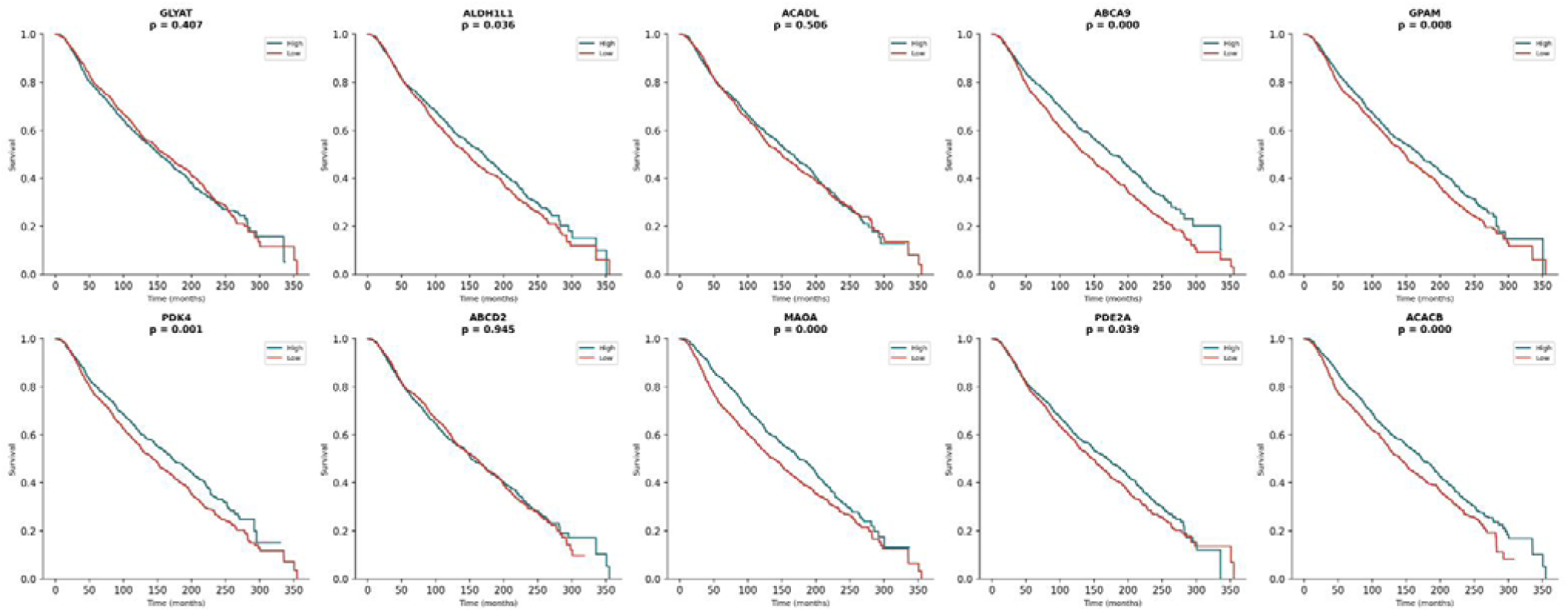
Kaplan–Meier overall survival — all 10 candidate genes (METABRIC). Kaplan–Meier overall survival curves for each of the 10 candidate genes individually in METABRIC (n = 1,980; median-dichotomised expression; log-rank p-values shown). Multiple genes reach significance in this larger, longer follow-up cohort. High expression: teal; low expression: red.

**Additional file 1: Figure S3.**
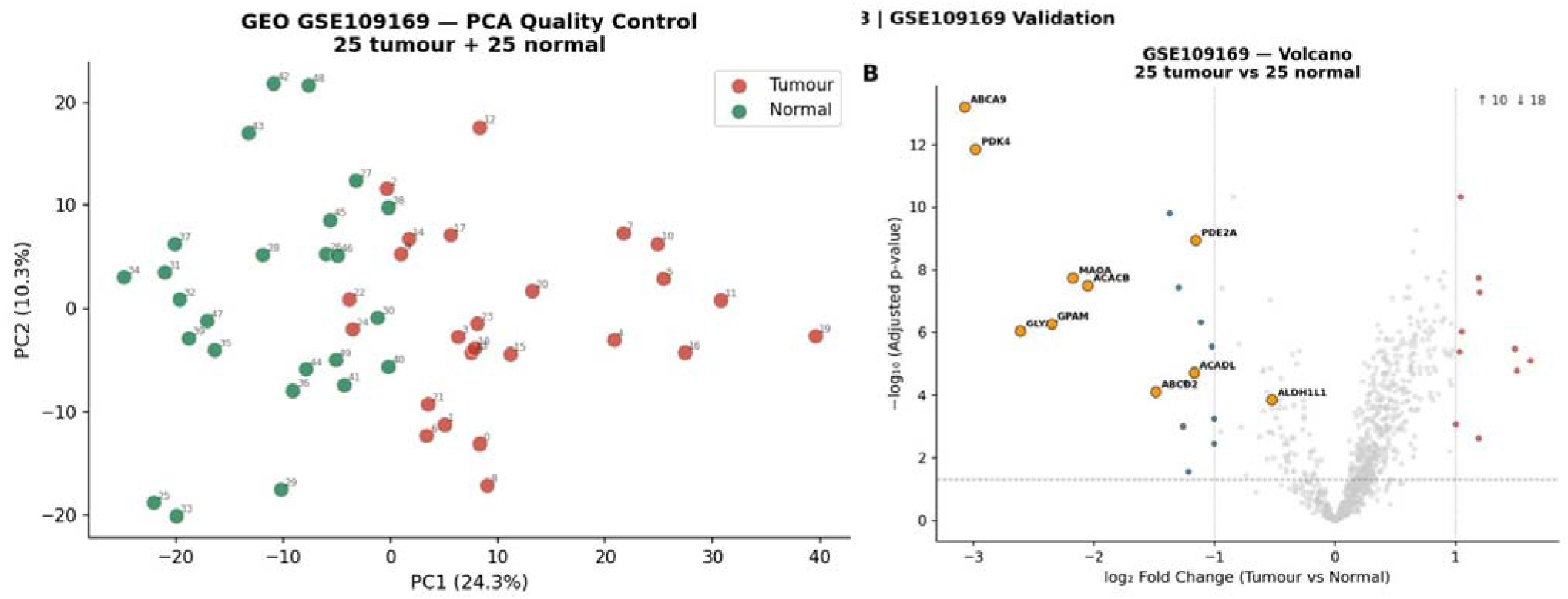
GSE109169 cross-platform validation. (a) PCA of GSE109169 samples (Affymetrix HuEx-1.0-ST; GPL5175; 25 matched tumour-normal pairs). PC1 (24.3% variance) shows partial separation between tumour (red) and normal (green) samples. (b) Volcano plot of differential expression (25 tumours vs 25 normals; Welch t-test; BH FDR). Orange circles indicate the 10 candidate genes, all individually replicated with concordant direction of effect. BH: Benjamini–Hochberg; FDR: false discovery rate; PCA: principal component analysis.

**Additional file 1: Figure S4.**
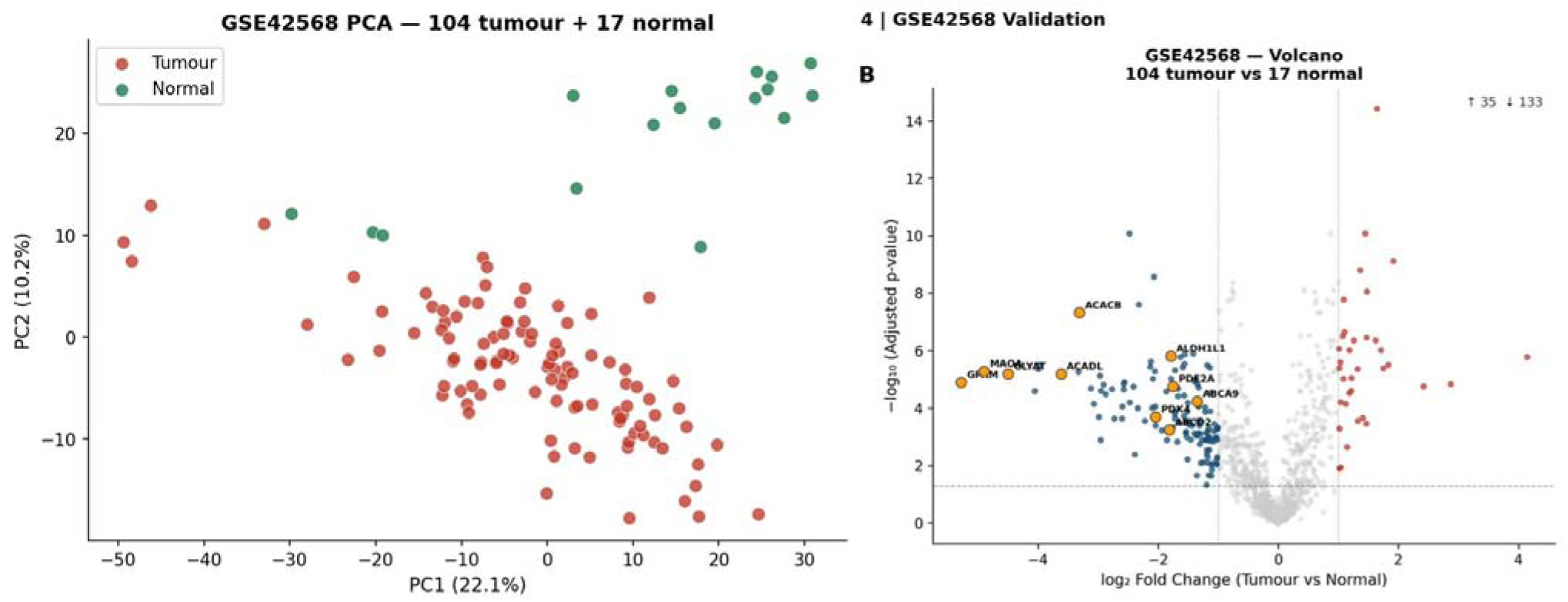
GSE42568 replication cohort validation. (a) PCA of GSE42568 samples (Affymetrix HG-U133 Plus 2.0; GPL570; 104 primary tumours, 17 normal-adjacent tissues). PC1 (22.1% variance) shows separation between tumour (red) and normal (green) samples. (b) Volcano plot of differential expression (104 tumours vs 17 normals; Welch t-test; BH FDR). Orange circles indicate the 10 candidate genes, all individually replicated. BH: Benjamini–Hochberg; FDR: false discovery rate; PCA: principal component analysis.

**Additional file 1: Figure S5.**
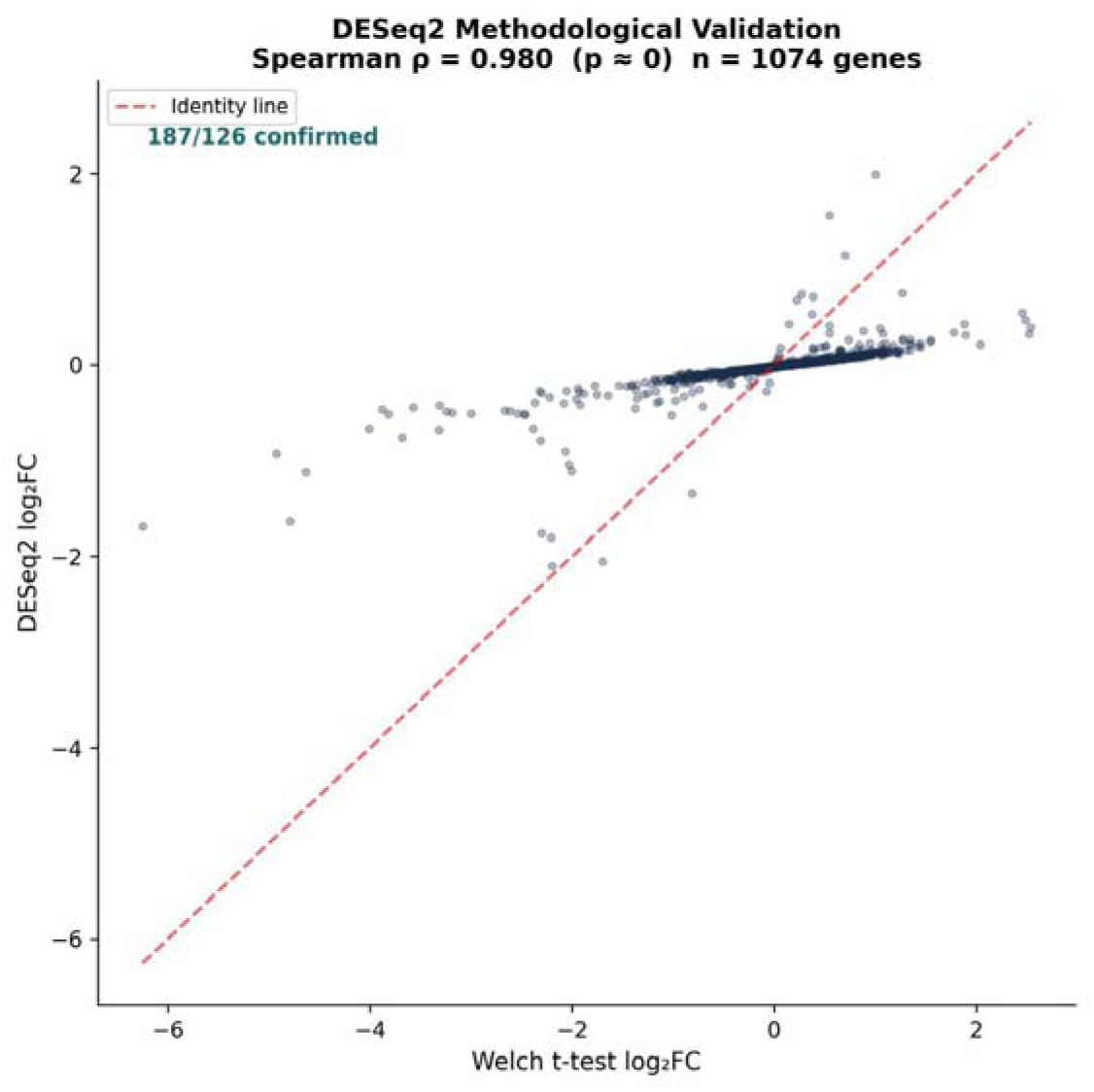
DESeq2 methodological cross-validation. Spearman rank correlation (ρ = 0.980; p ≈ 0; n = 1,074 genes) between DESeq2 and Welch t-test log₂ fold-change estimates across all testable MitoCarta 3.0 genes in TCGA-BRCA. Red dashed line: identity line. FC: fold change.

**Additional file 1: Figure S6.**
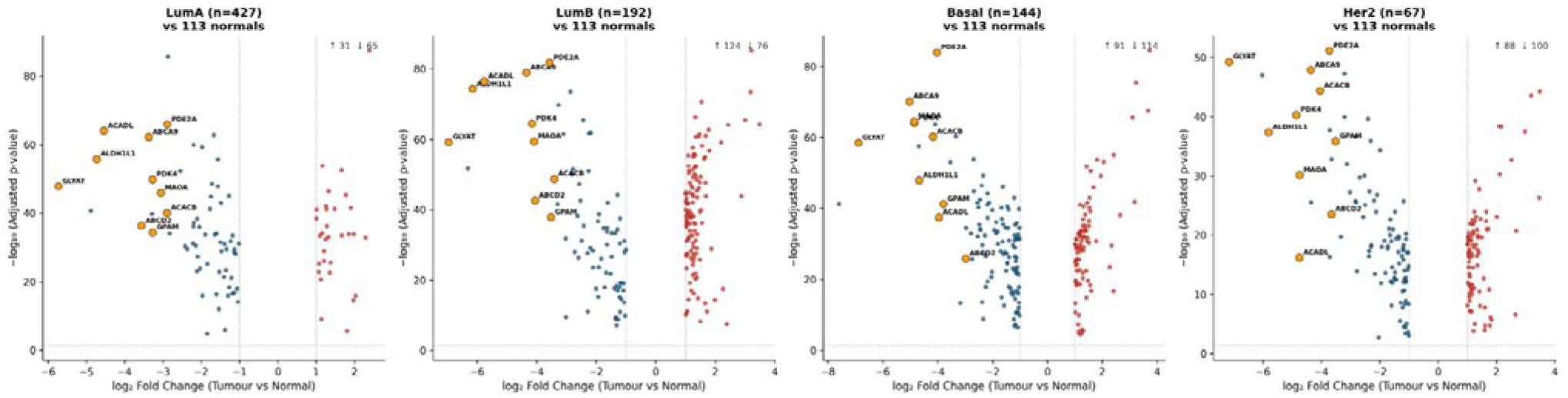
PAM50 subtype-specific volcano plots. Volcano plots of differential expression for each PAM50 subtype vs 113 normal-adjacent tissue samples: LumA (n = 427), LumB (n = 192), Basal (n = 144), HER2 (n = 67). Upregulated: red; downregulated: blue; non-significant: grey. Orange circles: 10 candidate genes, consistently downregulated across all subtypes. BH FDR thresholds applied. BH: Benjamini–Hochberg; FDR: false discovery rate.

## Notes

https://github.com/y7ktdd4cjd-lgtm/breast-cancer-mitochondrial-FAO

### Competing Interest Statement

The authors have declared no competing interest.

