## Additional File 2 for "Convergent suppression of nuclear-encoded mitochondrial fatty acid oxidation genes defines a pan-subtype signature in breast cancer: a multi-cohort transcriptomic study"

**REMARK Checklist — Reporting Recommendations for Tumor Marker Prognostic Studies**

Authors: Gomosani A, Marghalani H, Al Matar L | Reference: McShane LM et al. J Natl Cancer Inst. 2005;97(16):1180–1184. doi:10.1093/jnci/dji237

| **Item** | **REMARK guideline item** | **Response (this study)** | **Manuscript location** |
| --- | --- | --- | --- |
| **I. STUDY POPULATION** | | | |
| **1** | Describe the characteristics (e.g., disease stage or co-morbidities) of the study patients, including age and sex, and the laboratory characteristics of the tumor samples. | Primary discovery cohort: TCGA-BRCA, n = 1,219 samples (1,106 primary tumours, 113 normal-adjacent, 7 metastatic excluded). All female breast cancer patients. Age range, stage, and PAM50 subtype distribution detailed in Table 1. GDC dataset version 09-07-2024. External cohorts: GSE42568 (n = 121; 104 tumours, 17 normals; Affymetrix HG-U133 Plus 2.0); GSE109169 (n = 50; 25 matched pairs; Affymetrix HuEx-1.0-ST). Prognostic cohort: METABRIC (n = 1,980; Illumina HT-12 v3; 1,143 OS events; median follow-up ~10 years). | Methods — Data acquisition; Table 1 |
| **2** | Describe inclusion and exclusion criteria. | Inclusion: Primary tumour samples (GDC code 01) and normal-adjacent tissue (code 11). Exclusion: Metastatic samples (code 06; n = 7) excluded from all analyses. For external cohorts, all samples in GEO series matrices with available gene expression data were included. METABRIC samples with complete clinical annotation were used. | Methods — Data acquisition |
| **3** | Give the dates defining the period of accrual of patients into the study. | TCGA-BRCA data accessed: September 2024 (GDC v09-07-2024). Original TCGA-BRCA sample accrual dates are not specified in this analysis of publicly available data; refer to the TCGA BRCA marker paper (TCGA Network, Nature 2012). METABRIC: median follow-up ~10 years (Curtis et al. Nature 2012). GEO datasets: GSE42568 and GSE109169 (dates per GEO repository). | Methods — Data acquisition; Reference [25, 18] |
| **II. SPECIMEN CHARACTERISTICS** | | | |
| **4** | Describe the assay method used and provide (or reference) a detailed protocol for measuring the biomarker in the tumor specimens. | TCGA-BRCA: RNA-sequencing (RNA-seq), STAR aligner, FPKM-UQ normalisation. Processed data downloaded from GDC Data Portal. GSE42568: Affymetrix HG-U133 Plus 2.0 microarray (GPL570). GSE109169: Affymetrix HuEx-1.0-ST microarray (GPL5175). METABRIC: Illumina HT-12 v3 microarray. All expression data used as provided in public repositories. MitoCarta 3.0 used to define the nuclear-encoded mitochondrial gene set (1,136 genes; Rath et al. Nucleic Acids Res. 2021). | Methods — Data acquisition; External validation cohorts |
| **5** | Describe any handling of tumor material that could affect the quality of the data. | This study uses publicly available, uniformly processed data from established repositories (GDC, GEO, cBioPortal). Pre-analytical handling of original tissue samples was performed by the original data-generating consortia and is described in their respective primary publications. No additional processing was performed in this study beyond bioinformatic quality control (PCA, version harmonisation, metastatic sample exclusion). | Methods — Quality control |
| **III. ASSAY METHODS** | | | |
| **6** | Specify the number of samples assayed and the number of observations per patient. | TCGA-BRCA: 1,219 total samples (1,106 tumour + 113 normal; 7 metastatic excluded); one RNA-seq observation per sample. GSE42568: 121 samples (104 tumour + 17 normal); one microarray observation per sample. GSE109169: 50 samples (25 matched pairs); one microarray observation per sample. METABRIC: 1,980 samples; one observation per sample. | Methods — Data acquisition; Table 1 |
| **7** | Describe any quality control (QC) measures. | PCA was performed on all 1,219 TCGA-BRCA samples prior to any differential expression analysis to confirm tumour/normal separation, identify outliers, and detect batch structure. Ensembl gene ID harmonisation (version suffix stripping) ensured cross-dataset compatibility. Metastatic samples excluded before analysis. DESeq2 applied as independent methodological cross-validation (Spearman ρ = 0.980 vs Welch t-test). GEO cohort PCA performed to confirm sample separation. | Methods — Quality control; Methodological cross-validation; Additional file 1: Fig. S5 |
| **IV. STUDY DESIGN** | | | |
| **8** | State the hypothesis/objective clearly, including whether the study was designed before or after data collection. | Pre-specified hypothesis: Nuclear-encoded mitochondrial fatty acid oxidation (FAO) suppression constitutes a convergent, pan-subtype feature of breast cancer initiation. Candidate gene scoring formula was pre-specified and locked before any clinical testing or survival analysis, to prevent selection–outcome circularity. This is a retrospective analysis of publicly available data; the hypothesis was formulated prior to analysis. | Background; Methods — Candidate gene selection |
| **9** | State the primary and secondary endpoints. | Primary endpoint: Identification of a pan-subtype nuclear-encoded mitochondrial FAO suppression signature in TCGA-BRCA, confirmed by three-way intersection (all-stage ∩ Stage I ∩ pan-PAM50 DE). Secondary endpoints: (1) GSEA pathway enrichment; (2) Diagnostic performance (AUC) of candidate genes; (3) Clinical associations with PAM50, AJCC stage, T stage, nodal status, ER status, HER2 status, age; (4) Overall survival associations; (5) Independence from PIK3CA/TP53 mutation; (6) External replication in GSE42568 and GSE109169; (7) Prognostic validation in METABRIC. | Background; Methods (all subsections) |
| **10** | Clearly define what the marker analysis was intended to evaluate: prognostic factor, predictive factor, or surrogate endpoint. | This study evaluates the candidate genes and composite FAO suppression score as: (1) diagnostic markers (tumour vs normal discrimination; ROC/AUC); (2) prognostic markers (association with overall survival; KM and Cox analyses in TCGA-BRCA and METABRIC). The markers are not evaluated as predictive factors (i.e., predictors of response to specific therapy) in this study. | Methods — Diagnostic performance; Survival analysis |
| **V. STATISTICAL ANALYSIS METHODS** | | | |
| **11** | Describe all statistical methods, including those used to examine the association between the marker and patient outcome. | Differential expression: Welch two-sample t-test on log₂(FPKM-UQ+1) values, BH FDR correction. Significance: adjusted p < 0.05 and \|log₂FC\| > 1. Pathway enrichment: GSEA (pre-ranked by Welch t-statistic; MitoCarta 3.0 gene sets; FDR < 0.25). Replication: Hypergeometric test. Effect size concordance: Spearman rank correlation. Diagnostic performance: ROC-AUC (trapezoidal integration). Clinical associations: Chi-square (categorical) and Mann-Whitney U (age); BH FDR across 70 simultaneous tests. Survival: Kaplan-Meier log-rank; multivariable Cox regression (age, PAM50, ER status). Mutation analysis: Welch t-test, BH FDR across 20 simultaneous tests. Methodological validation: DESeq2; Spearman correlation of log₂FC estimates. | Methods (all statistical subsections) |
| **12** | If the study uses a subset of patients from a larger study, describe the rationale for focusing on this subset. | Not applicable. The full available TCGA-BRCA cohort was used for discovery. Stage I analysis used a pre-specified subgroup (n = 163 Stage I tumours) to assess whether dysregulation is detectable at disease initiation; this was a planned analysis, not a post-hoc subset selection. | Methods — Differential expression analysis |
| **13** | Describe how the marker was handled in the analyses; specify continuous analyses, cutoffs, and groupings. | For KM survival and clinical association analyses: candidate gene expression was median-dichotomised within the tumour sample population (high vs low), with the cutpoint determined as the median expression value across tumour samples — a pre-specified approach applied uniformly across all 10 genes. For Cox regression: the composite FAO suppression score was entered as a continuous variable. For ROC/AUC: individual gene expression used as continuous predictor against binary tumour/normal outcome. | Methods — Survival analysis; Clinical association analysis; Diagnostic performance |
| **14** | Explain how missing data were handled. | TCGA-BRCA GDC data: 100% clinical alignment rate across 1,219 samples (no missing clinical data). MitoCarta gene matching: 1,079/1,136 genes matched (95.0%; 57 unmatched genes absent from TCGA-BRCA expression data, not treated as missing). METABRIC: samples with complete clinical annotation for age, PAM50, and ER status were used in Cox regression; the total METABRIC n = 1,980 was used for KM analyses. No imputation was performed. | Methods — Data acquisition; External validation cohorts |
| **15** | Specify how statistical significance was determined. | Differential expression: BH FDR < 0.05 and \|log₂FC\| > 1 (both criteria required). GSEA: FDR < 0.25 per Subramanian et al. (2005) convention. Clinical associations: BH FDR < 0.05 across 70 simultaneous tests. Mutation analysis: BH FDR < 0.05 across 20 simultaneous tests. Survival: log-rank p < 0.05 (KM); adjusted p < 0.05 (Cox). External replication: hypergeometric p < 0.05 with 100% directional concordance. All multiple testing corrections applied using BH procedure. | Methods (all subsections) |
| **VI. RESULTS** | | | |
| **16** | State the number and characteristics of the patients studied. Describe the relationship between marker positivity and standard clinical and pathological variables. | TCGA-BRCA: 1,106 primary tumours and 113 normal-adjacent tissues. 966 tumours with PAM50 annotation. Clinical associations reported for all 10 candidate genes across PAM50, AJCC stage, T stage, nodal status, ER status, HER2 status, and age (Table 3). All 10 genes significantly associated with PAM50 subtype. ABCA9 was most broadly associated (6/7 clinical variables). FAO suppression score differed significantly by PAM50 subtype (ANOVA p = 1.54 × 10⁻²²). | Results — Dataset composition; Clinical associations; Table 1, 3; Fig. 5 |
| **17** | Report the results of the univariable analyses of the relationship between the marker and the study endpoint. | Kaplan-Meier analyses: ACADL (log-rank p = 0.040) and ABCD2 (p = 0.028) reached significance in TCGA-BRCA. Composite FAO suppression score in METABRIC: log-rank p = 7.82 × 10⁻⁷. Individual KM results for all 10 candidates in TCGA-BRCA and METABRIC are reported in Additional file 1: Figs. S1–S2. | Results — Survival analysis; Fig. 6; Additional file 1: Figs. S1–S2 |
| **18** | Report multivariable analyses if performed. Specify all variables entered in the model and their relationship to outcome. | Multivariable Cox proportional hazards regression in METABRIC, adjusted for age (continuous), PAM50 subtype (4 categories), and ER status (binary). MAOA: HR = 0.890 (95% CI 0.819–0.967; adjusted p = 0.009). Composite FAO suppression score: HR = 0.942 (95% CI 0.881–1.007; p = 0.078). The composite score did not reach significance after adjustment, interpreted as a directionally consistent trend. | Results — Survival analysis; Fig. 6c |
| **19** | If done, report results of any subgroup analyses that were performed. | PAM50-stratified DE analyses: LumA (n = 427), LumB (n = 192), Basal (n = 144), HER2 (n = 67), each vs 113 normals. Results in Additional file 1: Fig. S6 and Supplementary Figure 6. ANOVA of composite FAO score across PAM50 subtypes: p = 1.54 × 10⁻²² (Fig. 4b). Mutation status stratification (PIK3CA: 2/10 genes significant; TP53: 9/10 genes significant; Fig. 7). | Results — PAM50 section; Mutation analysis; Figs. 4b, 7; Additional file 1: Fig. S6 |
| **VII. DISCUSSION** | | | |
| **20** | Interpret the results in the context of the pre-specified hypothesis and other relevant data, and discuss any limitations. | Results are consistent with the pre-specified hypothesis that FAO suppression is a pan-subtype convergence point in breast cancer. FAO was the most significantly depleted MitoCarta pathway (NES = −2.130; FDR = 0.001), detectable at Stage I, with 100% directional concordance across two independent external cohorts. Limitations discussed explicitly: normal-adjacent tissue reference, transcriptomic vs functional FAO activity, observational design limiting causal inference, composite score Cox result not reaching significance, underpowering of GSE109169 for gene-level replication. Clinical relevance discussed in terms of pan-subtype therapeutic potential. | Discussion; Conclusions |

*Note: This checklist was completed in accordance with McShane LM, Altman DG, Sauerbrei W, Taube SE, Gion M, Clark GM. Reporting Recommendations for Tumor Marker Prognostic Studies (REMARK). J Natl Cancer Inst. 2005;97(16):1180–1184. doi:10.1093/jnci/dji237*
